# Dim blue light drives reversible fucoxanthin derivative accumulation in the pelagophyte *Pelagomonas calceolata*

**DOI:** 10.64898/2026.09.29.755320

**Authors:** Chloé Seyman, Laurie Bertrand, Aurélie Fossey-Jouenne, Céline Orvain, Alice Moussy, Benjamin Noel, Quentin Carradec, Adrien Thurotte

## Abstract

- Pigment composition and regulation are critical for efficient photosynthesis in the ocean, where light intensity decreases and the spectrum narrows with depth. Fucoxanthin (Fx), the main carotenoid of several microalgal lineages, harvests the blue-green light prevailing in the deep euphotic zone. In pelagophytes, the acylated derivative 19′-butanoyloxyfucoxanthin (19′-BFx) is abundant, yet its function and regulation remain unclear.
- Here, we investigated how irradiance and spectral quality shape photoacclimation in *Pelagomonas calceolata*, an abundant and cosmopolitan low-light pelagophyte. Cultures were grown under blue or white light across 2-60 µmol photons m⁻² s⁻¹. We analysed growth, PSII photophysiology, pigment composition, gene expression levels, and the impact of spectral-shifts on the 19’-BFx/Fx ratio.
- *Pelagomonas calceolata* grew optimally under dim blue light and displayed low non-photochemical quenching (NPQ) under assay conditions. Transcriptomes revealed broad remodelling driven mainly by irradiance and modulated by light colour, including carotenoid-related processes. Dim blue light progressively increased the 19′-BFx/Fx ratio, whereas white light kept it low and reversed the response after a blue-to-white shift.
- These results identify reversible, spectrum-dependent 19′-BFx accumulation as a key component of photoacclimation in *P. calceolata,* and separate irradiance-driven from spectrum-dependent cellular responses. This plasticity may explain the ecological success of *P. calceolata* in low-light oceans.

## Introduction

Phytoplankton contributes approximately half of the Earth’s primary production through photosynthesis in the euphotic zones of the oceans (Field *et al*., 1998; Mattei & Scardi, 2021). While light is required to support microalgal growth, excessive irradiance can induce photoinhibition, leading to reduced growth rate (Goss & Lepetit, 2015; Darvehei *et al*., 2018). At the ocean surface, microalgae are exposed to high-light conditions characterized by a broad-spectrum solar irradiance that includes ultraviolet (UV), visible, and near-infrared wavelengths. As depth increases, light intensity decreases and the spectral composition changes due to absorption and scattering of light by water molecules, coloured dissolved organic matter (CDOM), non-algal particles, and phytoplankton cells themselves (Jerlov, 1976). Short (< 400 nm) and long (> 550 nm) wavelengths are rapidly attenuated in the upper water column, whereas blue-green wavelengths (400-550 nm) penetrate deeper into the euphotic zone (Brunet *et al*., 2014; Hallmann, 2025). This vertical gradient in light intensity and quality creates distinct ecological niches that are occupied either by microalgal species genetically adapted to specific light environments or by species capable of acclimating to a broad range of light conditions (Kirkham *et al*., 2013; Brun *et al*., 2015; Jaubert *et al*., 2017). Characterizing the molecular mechanisms underlying adaptation and acclimation to these vertical light gradients is essential for understanding phytoplankton distribution in the oceans, particularly as climate-driven stratification of the upper ocean intensifies (Sallée *et al*., 2021; Hutchins & Tagliabue, 2024).

Algal lineages possess specific pigment repertoires that are regulated in response to variations in light intensity and spectral quality (Gong & Bassi, 2016; Hallmann, 2025). Red-lineage - derived algae, including ochrophytes, haptophytes, and many dinoflagellates, possess diverse carotenoid pigments, among which fucoxanthin (Fx) and its derivatives are typically the most abundant (Dautermann *et al*., 2020). Fx absorbs predominantly in the blue-green region of the spectrum, suggesting a key role in supporting photosynthesis in deeper oceanic layers where these wavelengths dominate (Wang *et al*., 2019). Its biosynthesis is regulated by both light intensity and spectral quality in several microalgae species (Wang *et al*., 2018; Yang & Wei, 2020; Zarekarizi *et al*., 2023). The regulation of carotenoid biosynthesis involves photoreceptor proteins, which are particularly diverse in red-lineage algae (Coesel *et al*., 2021). Aureochromes, cryptochromes, and rhodopsin photoreceptors absorb blue to green wavelengths, whereas phytochromes primarily absorb red and far-red light (Jaubert *et al*., 2017; Hallmann, 2025). This diversity of photoreceptors allows the microalgae to perceive different light environments and to adjust their cellular machinery or behavior through downstream signaling pathways (Jaubert *et al*., 2017; Coesel *et al*., 2021; Coesel, 2024). Notably, heterologous expression of a cryptochrome gene has been reported to correlate with fucoxanthin accumulation in *Phaeodactylum tricornutum*, suggesting a regulatory role in carotenoid biosynthesis (Zhang *et al*., 2022).

Carotenoid compositional remodelling in microalgae represent an important component of photoacclimation, as carotenoids participate both in light harvesting and in the regulation of excitation-energy dissipation (Dubinsky & Stambler, 2009). Fx acyloxy derivatives 19′-hexanoyloxyfucoxanthin (19′-HFx) and 19′-butanoyloxyfucoxanthin (19′-BFx) have been reported in haptophytes and pelagophytes (Sanz *et al*., 2015). Using pigments isolated from the haptophyte *Emiliania huxleyi* and the pelagophyte *Aureococcus anophagefferens,* ultrafast spectroscopy has shown that acylation of fucoxanthin alters its excited-state dynamics, modulating light harvesting and potentially contributing to photoacclimation or photoprotection (Staleva-Musto *et al*., 2018). In *E*. *huxleyi*, 19′-HFx abundance increases under low irradiance, particularly under blue light (Garrido *et al*., 2016). Different species of *Phaeocystis* produce either 19′-HFx, 19′-BFx, or both pigments, with relative proportions varying according to environmental conditions and geographic location (Zapata *et al*., 2004; Van Leeuwe *et al*., 2014; Wang *et al*., 2022). These observations highlight both the molecular diversity of haptophyte pigments and the fine-tuned regulation of pigment composition in response to the environment. Among pelagophytes, a diverse class of microalgae encompassing open ocean and coastal species, 19′-BFx is generally the dominant Fx derivative and has long been used as a chemotaxonomic marker of this lineage (Bjørnland *et al*., 1989; Andersen *et al*., 1993; Dimier *et al*., 2009). In the bloom-forming pelagophyte *A. anophagefferens*, the 19′-BFx/Fx ratio increases under high-light conditions, and 19′-BFx accumulates more strongly during the stationary growth phase (Alami *et al*., 2012; Cui *et al*., 2025). Its accumulation is concomitant with a reduced antenna efficiency, suggesting a photoprotective role for 19′-BFx (Alami *et al*., 2012). Despite their abundance and dynamic regulation patterns, the physiological functions of Fx derivatives remain poorly understood.

The pelagophyte *Pelagomonas calceolata* is a cosmopolitan species and one of the most abundant phytoplankton taxa in the open ocean (Andersen *et al*., 1993; Worden *et al*., 2012; Guérin *et al*., 2022). It is typically more abundant in the deep chlorophyll maximum (DCM) than in surface waters and is often found deeper in the water column than many other phytoplankton lineages (Cabello *et al*., 2016; Latasa *et al*., 2016; Guérin *et al*., 2022). Both field observations and culture experiments have demonstrated that *P. calceolata* exhibits remarkable acclimation capacities to low-light, which may partly explain its ecological success in this environment (Timmermans *et al*., 2005; Kang *et al*., 2021; Coale *et al*., 2026).

In this study, we investigated photoacclimation in the pelagophyte *P. calceolata* by cultivating cells under different intensities of blue and white light, used as proxies for the light conditions encountered across the euphotic zone. By combining physiological measurements, pigment profiling and transcriptomics we characterize the coordinated physiological and molecular responses underlying photoacclimation in this abundant and cosmopolitan pelagophyte, and provide new insights into the regulation of fucoxanthin-derived pigments such as the poorly understood 19′-BFx.

## Material and methods

### Culture and growth monitoring of P. calceolata

*P. calceolata* strain RCC100, from the Roscoff Culture Collection, was grown in artificial seawater (ASW) supplemented with L1 medium (Bigelow L1 Medium Kit), in 150 mL Erlenmeyer flasks under 14:10 light:dark photoperiod. The ASW L1 medium was prepared following the protocol described in Guérin et al. 2025. For the first light experiment, 15 mL cultures were grown under 14 intensities of blue (max emission: 455 nm) or white LEDs between 0 and 70 µmol photons m^−2^ s^−1^, measured with a LI-COR LI-180 portable spectrometer (emission spectra presented in Supporting Information Fig. S1). The dark condition (0 µmol photon m^−2^ s^−1^) was used as a negative control for growth and fluorescence. For the transcriptomic and pigment analyses, 60 mL cultures were exposed to blue (BL) or white (WL) light at four intensities (4, 10, 20 and 60 µmol photons m^−2^ s^−1^). The cultures were carried out in triplicate (0 and 60 µmol photons m^−2^ s^−1^ conditions) or quadruplicate (4, 10 and 20 µmol photons m^−2^ s^−1^ conditions). The cell concentration at the start of the experiment was set at 500,000 cells mL^−1^. For the monitoring experiment of 19′-BFx accumulation, triplicate cultures of *P. calceolata* were initially exposed to 4 µmol photons m^−2^ s^−1^ of white light for one week. The cultures were then transferred to 4 µmol photons m^−2^ s^−1^ of blue light for one week. After 7 days, the cultures were split into two groups: one was switched back to white light, the other was kept under blue light for an additional week. In parallel, a control culture was kept under white light for 3 weeks. Cell growth was monitored throughout all experiments by *in vivo* fluorescence (excitation: 470 nm; emission: 665-720 nm, Qubit™ 3 Fluorometer, Invitrogen) and cell counting by flow cytometry (CytoFLEX, Beckman Coulter), applying a threshold based on side scatter (SSC) and B690 fluorescence. Measurements were performed daily for the 19′-BFx experiment and every 2-3 days for the light intensity experiments.

Specific growth rates (µ) were estimated by linear regression of ln(cell abundance) against time using sliding windows of six consecutive time points, retaining the highest positive µ for each growth curve. Growth-irradiance relationships were described using the Peeters rational photoinhibition model (Peeters & Eilers, 1978), corresponding to model Ph02 implemented in the R package *piCurve* (Amirian & Irwin, 2025).

### PAM measurement

Chlorophyll *a* fluorescence was measured using a Dual-Modulation Kinetic Fluorometer FL 6000 (Photon Systems Instruments) at 20°C. Measurements were performed during the exponential phase in biological triplicates. Samples were dark-adapted for 5 minutes before measurement. Measurements were performed at 623 nm using the Quenching analysis program provided by Photon Systems Instruments. The maximum quantum yield of photosystem II (Fv/Fm) was calculated as (Fm-F_0_)/Fm, where F_0_ is the minimum fluorescence and Fm is the maximum fluorescence of dark-adapted cells. Non-Photochemical Quenching (NPQ) was calculated using the last saturation pulse (Fm’) under actinic light and applying the (Fm-Fm’)/Fm’ formula.

### Pigments extraction and quantification

For pigment analysis across the blue- and white-light irradiance series, a 200-µL aliquot was collected from 3 biological replicates of each condition during the exponential phase. For the 19′-BFx/Fx ratio monitoring experiment, 1 mL samples were taken from each flask 6 hours after the beginning of every phase (t0+6h and t7+6h) and then every day at the same hour (10 a.m). The samples were centrifuged for 10 minutes at 10,000 rcf, and the pellets were flash-frozen in liquid nitrogen before being stored at −80°C. Cell pellets were resuspended in 10 mM TRIS HCl (pH 7.8) in 100 µL of methanol:acetone (70:30), and incubated for 10 minutes with periodic agitation. The samples were then centrifuged at 10,000 rcf for 5 minutes, and the supernatant was collected and filtered through a 0.22 µm PVDF membrane to remove cell debris.

Pigment extracts were separated on an ACQUITY UPLC BEH C18 column (100 × 2.1 mm; 1.7 µm; Waters) fitted to an Ultimate 3000 UHPLC RS 1034 bar system equipped with a UV-Vis detector (Thermo Fisher Scientific), with the column oven set to 40°C. The mobile phase consisted of solvent A (10 mM ammonium bicarbonate, adjusted to pH 7) and solvent B (methanol:acetonitrile, 7:3, v/v), delivered at 0.25 mL min⁻¹. The injection volume was 5 µL. Pigments were eluted over 20 min with a linear gradient from 50% to 95% B. Pigments were detected by UV-Vis absorbance at 443 nm, and their identification was carried out using pigment standards of chlorophylls *a* and *c3*, diadinoxanthin, diatoxanthin and 19’-BFx as well as a photosynthetic pigment mix (DHI™ Laboratory Products). Chromatograms presented in this paper were plotted with R and baseline was corrected with *baseline* R package *(v. 1.3-5)*. Pigment quantification was based on the area under the curve of each chromatogram peak, normalized to the cell concentration at the time of sampling. Statistical tests were performed using a one-way ANOVA, followed by a Tukey test, using the TukeyHSD function in R. Inter-replicate variability due to UHPLC injection was accounted for by including replicate as a blocking factor in the ANOVA model (Value ∼ Condition + Replicate). For data visualization only, batch effects were removed using the removeBatchEffect function from the *limma* R package (Ritchie *et al*., 2015).

### RNA extraction and sequencing

Cultures were harvested during the exponential phase at cell concentrations between 2.4 × 10 and 3.6 × 10 cells mL⁻¹. Cultures were filtered on a 1.2 µm Isopore filter (Polycarbonate membrane, 47 mm diameter), on a glass filtration unit connected to a peristaltic pump. Filters were immediately transferred to a 15 mL Falcon tube, flash-frozen in liquid nitrogen, and stored at −80°C until RNA extraction. Total RNA from *P. calceolata* was extracted using the RNeasy Plus Universal Mini Kit (Qiagen). Extracts were then digested with 4 U of TURBO™ DNase (2 U/µL) (Thermo Fisher Scientific) and subsequently purified using the RNA Clean & Concentrator-5 kit (Zymo Research). RNA-seq libraries were generated from 500 ng of purified RNA using the Illumina Stranded mRNA Prep, Ligation kit, following the standard protocol described in Guérin et al. 2025. The librairies were amplified using 15 PCR cycles, and sequenced on an Illumina NovaSeq X Plus sequencer (Illumina, San Diego, CA, USA) in 2×151 bp, targeting 30 million read pairs. Raw reads were first trimmed to remove Illumina adaptors and primer sequences, and low-quality bases (Qb < b20) were clipped from both read ends. Reads were further truncated at the position of the second ambiguous nucleotide (N), and any read shorter than 30 nt after trimming was discarded, using a custom adaptation of the fastx_clean tool (Warner, 2026). Finally, residual rRNA reads were removed using SortMeRNA v2.1 against the SILVA database (Kopylova *et al*., 2012; Quast *et al*., 2013).

### Transcriptomic analyses

RNA-seq reads were aligned to the gene models of *P. calceolata* genome (strain RCC100) using bwa-mem2 version 2.2.1. Aligned reads with a minimum length of 50 bp, an identity above 95% and at least 80% of their length were retained. Nuclear genes covered with at least 10 reads across all samples were kept for downstream analysis. Differential gene expression analyses between light conditions were performed on raw counts using the *DESeq2* R package (Love *et al*., 2014), with the 4 µmol photons m⁻² s⁻¹ condition set as the reference level for light intensity comparisons. A threshold of adjusted p-value < 0.01 and |log₂FC| ≥ 1 was used to define differentially expressed genes for gene-level analyses. For the Principal Component Analysis (PCA) and the Pearson correlation plot, raw counts were normalized to stabilize variance using the varianceStabilizingTransformation (vst) function from *DESeq2*. The PCA was performed on the 500 genes with the highest variance using the plotPCA function from the *DESeq2* package. A Uniform Manifold Approximation and Projection (UMAP) was computed on normalized read counts aligned on all *P. calceolata* genes using the R package *umap* (McInnes *et al*., 2020). Genes were coloured according to their differential expression status, distinguishing genes differentially expressed in at least one of the 3 light intensity contrasts (4 vs 10, 4 vs 20 and 4 vs 60 µmol photons m⁻² s⁻¹) under BL and WL, and genes differentially expressed between WL and BL at 4, 10, 20, or 60 µmol photons m⁻² s⁻¹.

### Gene set enrichment analysis

To identify functions enriched under low-light and high-light conditions, enrichment analyses based on Pfam domains and Gene Ontology (GO) annotations were performed. An over-representation analysis was conducted using a hypergeometric test implemented in the *DiCoExpress* pipeline (Lambert *et al*., 2020), with p-values adjusted for multiple testing using the Benjamini-Hochberg method. Enrichment analyses were performed on sets of DEGs defined using an adjusted p-value threshold of 0.01 and a relaxed log₂FC cutoff (|log₂FC| ≥ 0.585, corresponding to a fold-change ≥ 1.5).

### Identification of genes involved in pigment biosynthesis, photosystem assembly and photoreception

Nuclear genes putatively involved in photoreception, pigment biosynthesis, and photosystem assembly and stability were identified based on the functional annotation of *P. calceolata* genes (Supporting Information Table S4). LOV-domain-containing proteins were identified using *hmmer* tool version 3.4 performed with reference sequences from Coesel et al. (2021). Fourteen LOV-domain-containing genes were detected, including eight aureochromes and one phytochrome-like proteins. Aureochromes were identified based on the co-occurrence of PAS/LOV (IPR000014) and bZIP domains (IPR004827 or cd14809). The phytochrome-like protein was detected based on the presence of the PHY domain (IPR001294). Phytochrome-like proteins closely related to that of *P. calceolata* were identified using Blastp on the NCBI blast server against nr databases (BLOSUM62, default parameters) (Altschul *et al*., 1990), and on UniProt (The UniProt Consortium, 2025). The position of functional domains were determined using InterProScan 6 (Blum *et al*., 2026), and 3D protein structures were generated using AlphaFold 3 (Abramson *et al*., 2024). The seven cryptochromes were identified through the presence of domains annotated as “cryptochrome/DNA photolyase” (IPR002081, IPR036155, IPR036134). Rhodopsins were identified from annotations containing significant match with the Pfam domain “rhodopsin” (PF10192, PF18761, PF01036). Rhodopsins were assigned to families by multiple protein sequence alignment against a subset of 116 rhodopsin proteins representing different groups (enzyme rhodopsins, channelrhodopsins, sensory rhodopsins, ion-pump rhodopsins and heliorhodopsins) from Coesel *et al*., (2021) using MUSCLE (Edgar, 2004). The alignment was then trimmed with goalign version 0.3.1 (Lemoine & Gascuel, 2021) to remove sites containing more than 80% gaps. A phylogenetic tree was constructed with IQ-TREE (version 3.0.1; Wong *et al*., 2026) using 100 non-parametric bootstrap replicates (VT+F+R6 model chosen by ModelFinder). Branch support was then assessed by Transfer Bootstrap Expectation (TBE, Lemoine *et al*., 2018) using the Booster tool available here: https://booster.pasteur.fr/. The tree was subsequently annotated and coloured with iTOL (Letunic & Bork, 2024).

Genes involved in chlorophyll and carotenoid biosynthesis were identified using IPR, Pfam, and KOfam annotations on *P. calceolata* genes, using either gene names (e.g., ChlD, ChlM) or functional annotations (e.g., “protochlorophyllide reductase”, “zeaxanthin epoxidase”). The same annotation-based approach was applied to identify nuclear genes involved in photosystem assembly and stability.

## Results

### Growth and photosynthetic efficiency under different light conditions

To investigate the growth capacity of *P. calceolata* under different lights, we exposed cultures to 14 different intensities of blue light (BL) or white light (WL), between 2 and 70 µmol photons m^−2^ s^−1^. The optimal light intensity for *P. calceolata* growth differed between light conditions, with a lower optimum under blue light than under white light (10.8 vs 17.9 µmol photons m⁻² s⁻¹, respectively; Fig. 1a), with maximum growth rates of 0.360 and 0.387 d⁻¹, respectively. Model fits were strong under both conditions (adjusted R² = 0.94 and 0.98, respectively). At low irradiance, cultures grew slowly but eventually reached relatively high cell densities (Supporting Information Fig. S2). At higher irradiance, cultures showed faster initial growth but an earlier decline, consistent with the onset of photoinhibition, which limited biomass accumulation despite relatively high growth rates reached. These results indicate that *P. calceolata* is well adapted to low irradiance but subject to photoinhibition at higher light intensities.

**Fig. 1.**
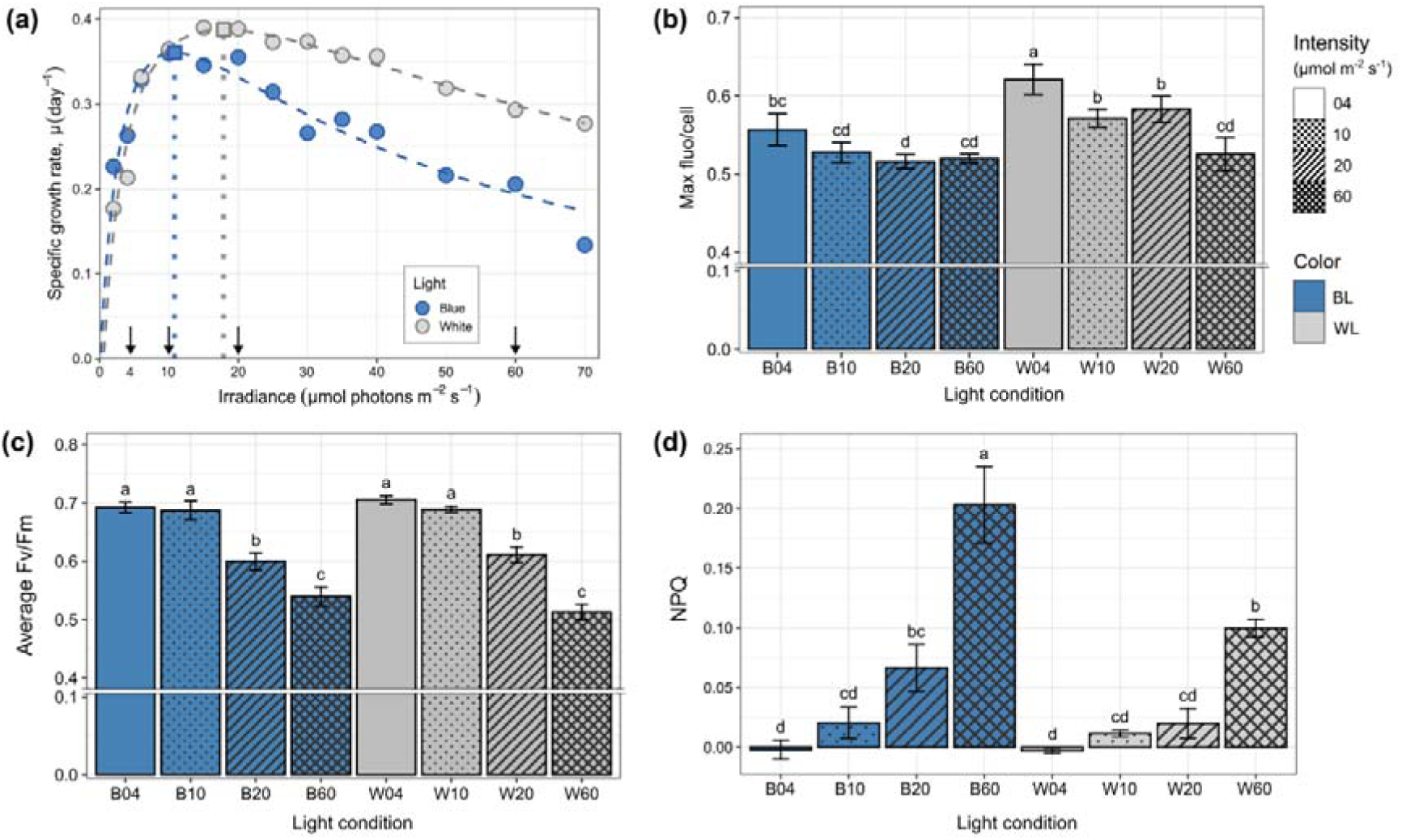
*P. calceolata* growth, fluorescence per cell, photosynthetic efficiency and photoprotection. **(a)** Specific growth rate (µ) according to irradiance under blue and white lights. Dashed curves represent fits of the Peeters rational photoinhibition model, and vertical dotted lines indicate the estimated optimal irradiance (Iopt). Arrows on the x axis represent the conditions selected to perform physiological and molecular analyses. **(b)** Maximum fluorescence per cell, over 35 days of culture. **(c)** Photosynthetic efficiency (Fv/Fm) and **(d)** NPQ of *P. calceolata* cultures in the 4 light conditions, measured the day of cell harvesting for transcriptomic analyses (between 11 and 25 days of culture). Error bars in panels b-d represent the standard deviation, and significant differences between conditions are indicated by letters on top of each bar (one-way analysis of variance (ANOVA) followed by Tukey’s multiple comparison).

We selected four irradiances (4, 10, 20 and 60 µmol photons m⁻² s⁻¹) to repeat the experiment under replicated conditions and characterize the physiological responses of *P. calceolata* (Supporting Information Fig. S3). Chlorophyll (Chl) fluorescence per cell was higher under low irradiance, particularly under WL (Fig. 1b). Photosynthetic performance was assessed by measuring the maximum quantum yield of photosystem II (Fv/Fm) during exponential growth. The highest Fv/Fm values were measured at 4 µmol photons m⁻² s⁻¹ under both BL and WL, and Fv/Fm declined progressively with increasing growth irradiance (Fig. 1c). Endpoint nonphotochemical quenching (NPQ), calculated using the final saturation pulse applied under actinic light, increased with growth irradiance under both BL and WL (Fig. 1d). Nevertheless, *P. calceolata* displayed low NPQ under the assay conditions, accompanied by declining Fv/Fm with increasing irradiance.

### Pigment composition and abundances

To assess the impact of light intensity and spectral quality on pigment composition, we quantified photosynthetic pigments of *P. calceolata* using Ultra High Performance Liquid Chromatography (UHPLC). Normalizing pigment abundance by cell concentration revealed that all pigments, except Diatoxanthin (Dt), decreased with increasing light intensity, under both BL or WL (Fig 2a, Supporting Information Fig. S4). This pattern was consistent with the reduction of photosynthetic efficiency observed at higher irradiance (Fig. 1c). To better resolve compositional changes, pigment abundances were therefore normalised to chlorophyll *a* abundance.

**Fig. 2.**
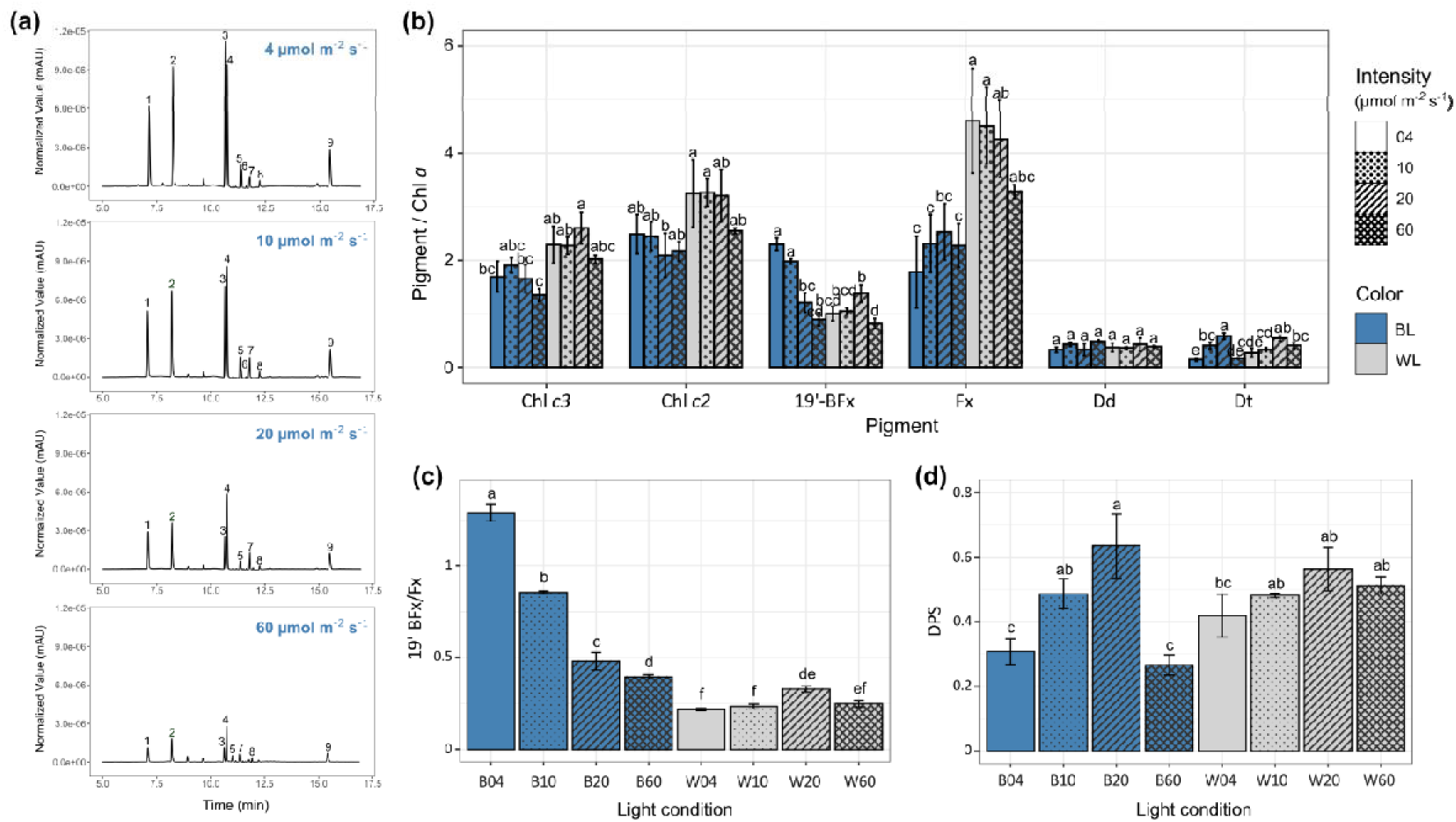
Variation in pigment abundances under BL or WL at different intensities. **(a)** UHPLC chromatograms at 443 nm of pigments from *P. calceolata* cultivated under BL at 4, 10, 20 or 60 µmol photons m^−2^ s^−1^ (top to bottom). Peak heights were normalized to cell concentration. Pigments were identified as follows: 1 = Chlorophyll *c2*, 2 = Chlorophyll *c3*, 3 = 19’-Butanoyloxyfucoxanthin (19′-BFx), 4 = Fucoxanthin (Fx), 5 = Diadinoxanthin (Dd), 6 = Dinoxanthin, 7 = Diatoxanthin (Dt), 8 = Zeaxanthin (putative), 9 = Chlorophyll *a*. **(b)** Pigment abundance ratios in *P. calceolata* relative to Chl *a*. Light intensities (µmol photons m^−2^ s^−1^) are indicated by different hatching patterns for BL and WL conditions. **(c)** Average of 19′-BFx/Fx ratio and **(d)** average of DPS ratio (Dt/[Dt+Dd]) under different intensities of blue (B) and white (W) lights. For all panels, error bars indicate the standard deviation and significant differences between conditions are indicated by different letters on top of each bar (one-way analysis of variance (ANOVA) followed by Tukey’s multiple comparison).

The most striking observation was the inversion of peak heights between 19’-Butanoyloxyfucoxanthin (19′-BFx) and fucoxanthin (Fx) abundances with light intensity under blue light (Fig. 2a). The relative abundance of 19’-BFx decreased with increasing light intensities under BL, but remained low across all light intensities under WL. In contrast, the relative abundance of Fx did not vary significantly with light intensity, but was consistently higher under WL than under BL (Fig. 2b). The relative abundances of Chl *c3* and *c2* showed no significant variation with light intensity, although both were slightly more abundant under WL than BL. Considering 19′-BFx as a derivative of Fx, we compared the 19′-BFx/Fx ratios across the different light conditions (Fig. 2c). The ratio was highest under very low BL (4 µmol photons m^−2^ s^−1^), with 19′-BFx being more abundant than Fx, and decreased sharply with increasing light intensity. By contrast, under WL, the ratio was lower than under BL and did not vary with light intensity, except at 20 µmol photons m^−2^ s^−1^.

Finally, the xanthophyll-cycle pigments diadinoxanthin (Dd) and diatoxanthin (Dt) showed the lowest relative abundance to Chl *a* among all pigments quantified (Fig. 2b). The relative abundance of the photoprotective pigment Dt increased progressively with irradiance under both BL and WL conditions whereas the relative concentration Dd pigment exhibited the most stable pattern, with its relative abundance remaining remarkably consistent across all light conditions tested. Accordingly, the de epoxidation state (DPS, Dt/(Dd+Dt)), which reflects the photoprotective conversion of Dd to Dt within the xanthophyll cycle, increased with light intensity accordingly (Fig. 2d) and closely paralleled with the NPQ levels measured under different light conditions (Fig. 1d). Under BL, DPS increased with light intensity and reached a maximum at 20 µmol photons m^−2^ s^−1^, before decreasing at the highest irradiance (60 µmol photons m^−2^ s^−1^). Although this observation could reflect a limitation of the de-epoxidation process beyond a certain high-light threshold, we cannot exclude that it instead stems from the uncertainty associated with quantifying the low abundance of Dt and Dd in high-light cultures.

### 19′-BFx/Fx ratio dynamics following spectral shifts

To investigate the kinetics or changes of 19’-BFx/Fx ratio following a shift in light quality, *P. calceolata* cells were first acclimated to low WL for one week, then exposed to low BL for 14 days or maintained at low WL (Fig. 3a,b). The irradiance was maintained at 4 µmol photons m⁻² s⁻¹ for both light colors during the entire experiment. During the 14 days under low BL, we observed a gradual increase of the 19’-BFx/Fx ratio, even at days 13 and 14 where cultures have reached their maximal concentration (Fig. 3a,b). Starting from an initial ratio of 0.5 (Fx twice more abundant than 19′-BFx), the balance shifted in favor of 19′-BFx after 9 days, reaching values above 1.2. In contrast, during the 14 days under dim WL, the ratio slightly decreased, from 0.7 to 0.5. Thus, dim blue light progressively increased the 19′-BFx/Fx ratio, whereas returning cultures to white light initiated a decline in this ratio.

**Fig 3.**
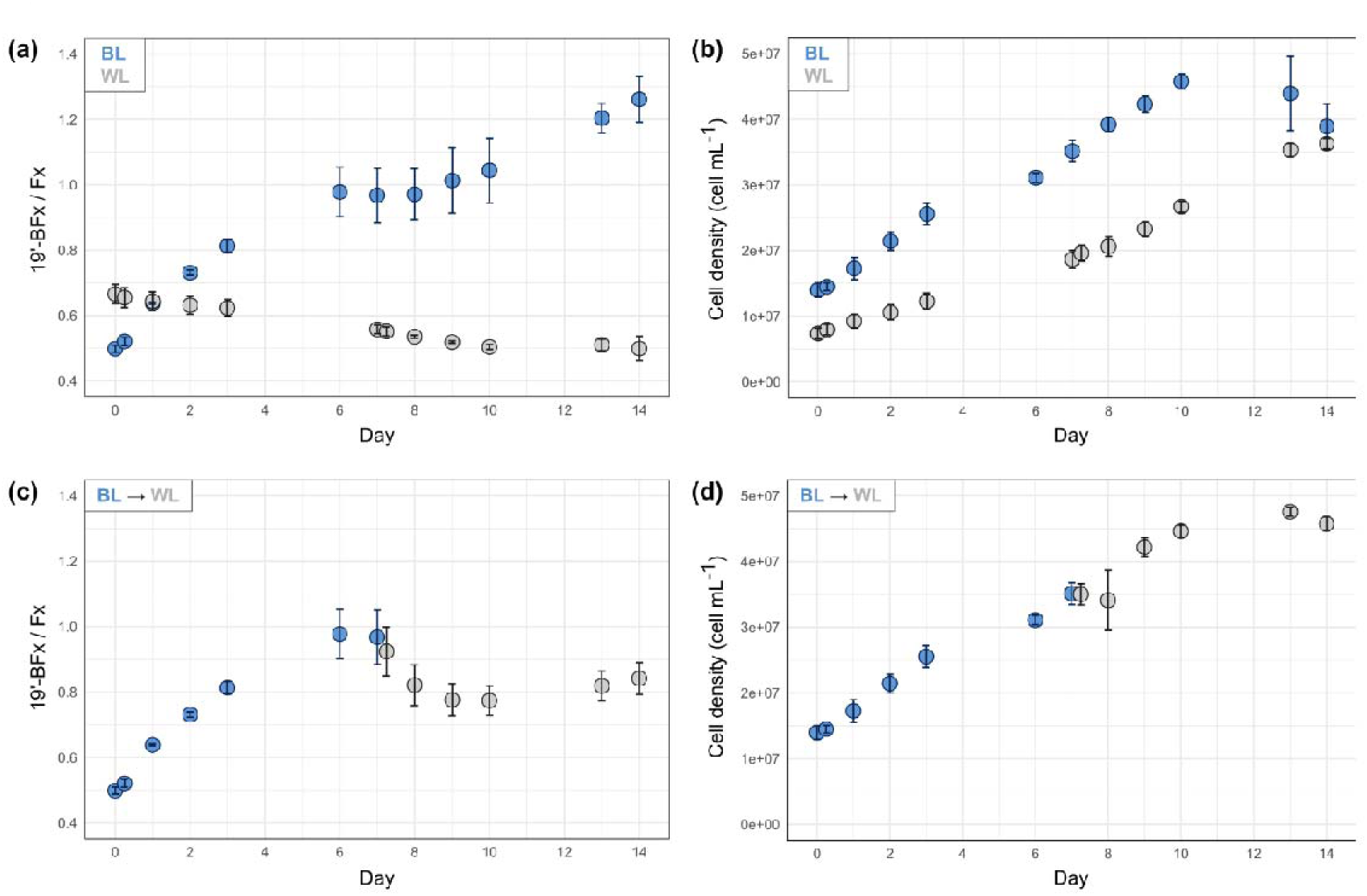
Dynamics of the 19’-BFx/Fx ratio and *P. calceolata* growth in response to alternating BL and WL conditions. Evolution of 19’-BFx/Fx pigment ratio **(a,c)** and *P. calceolata* cell growth **(b,d)** under 14 days of BL or WL (4 µmol photons m^−2^ s^−1^) **(a,b)**, and under 7 days of BL followed by 7 days of WL **(c,d)**. All cultures were initially acclimated to WL for one week. Blue dots correspond to cultures under BL, and grey dots correspond to cultures under WL. Error bars represent standard deviation between triplicates.

In parallel, triplicate cultures underwent the same initial acclimation under low WL, followed by 7 days under low BL and a further 7 days under low WL (Fig. 3c,d). Following the transfer from BL to WL, a decrease in the 19′-BFx/Fx ratio was detected at the first sampling point, 6 h after transfer (Fig. 3c). The spectral shift did not significantly affect cell abundance over the course of the experiment (Fig. 3d).

### Global patterns of transcriptomic regulation induced by light changes

To understand how *P. calceolata* acclimates to varying light intensities and spectral qualities at the gene expression levels, we performed a transcriptomic analysis on mRNAs extracted from the cultures acclimated to 4, 10, 20 and 60 µmol photons m^−2^ s^−1^ under BL or WL. A correlogram confirmed that samples clustered consistently by condition based on their expression levels, with samples at 60 µmol photons m^−2^ s^−1^ diverging most strongly from the others (Supporting Information Fig. S5). The Principal Component Analysis (PCA) on normalised read counts clearly separated samples according to light intensity on the first axis (70% of the variance of the top 500 genes) and BL from WL on the second axis (11% of the variance) (Fig. 4a). In the same way, the heatmap of normalized expression of genes with the highest variance shows a sample clustering that groups first by intensity (4-10 µmol photons m^−2^ s^−1^ and 20-60 µmol photons m^−2^ s^−1^), and then by color (Supporting Information Fig. S6). These observations indicate that the light intensity exerts a stronger influence on *P. calceolata* gene expression than the light color.

**Fig. 4.**
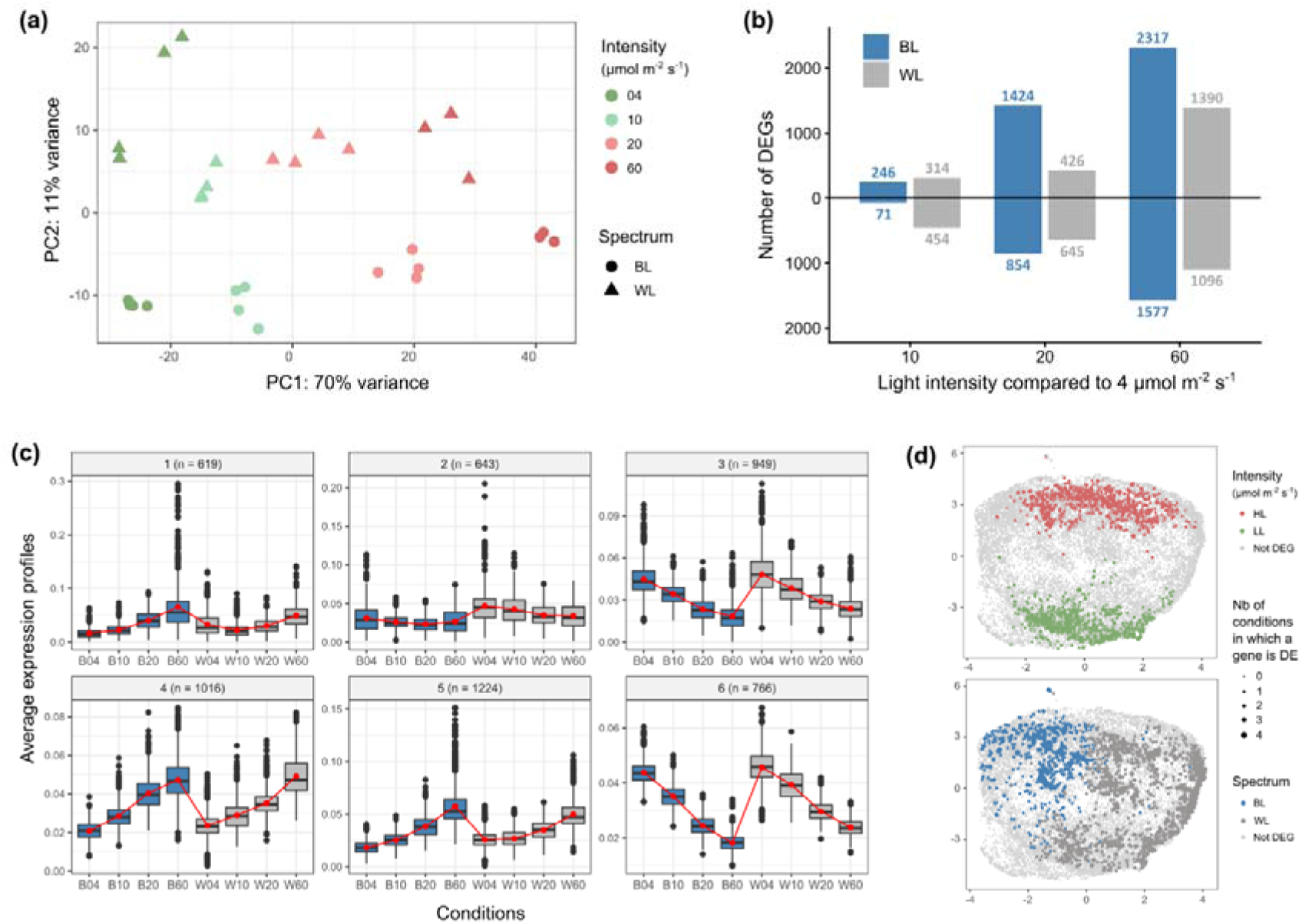
Global patterns of gene expression levels at different light intensities under BL or WL. **(a)** Principal Component Analysis (PCA) of the 500 genes with the highest variance across all conditions. The color of the dots represents the light intensity and the shape represents the light color. Dots with identical colors and shapes are biological replicates. **(b)** Number of upregulated genes (top) or downregulated genes (bottom) at 10, 20 and 60 µmol photons m^−2^ s^−1^ compared to 4 µmol photons m^−2^ s^−1^ (adjusted p-value < 0.01). **(c)** UMAP of gene expression levels profiles across all light conditions. DEGs with a |log₂FC| ≥ 1 between low and high light (up) and between blue and white light (down) are colored. Dot sizes reflect the number of conditions where each gene is differentially expressed. **(d)** Co-expression profiles of DEGs with |Log₂FC| ≥ 1 from the 30 *P. calceolata* samples grown under four light intensities of blue light (B) and white light (W).

We identified a total of 4,218 and 2,989 DEGs according to light intensity variations under BL and WL respectively (Fig. 4b, Supporting Information Table S3). The number of DEGs increased with the magnitude of the intensity difference from 317 DEGs between 4 and 10 µmol photons m^−2^ s^−1^ to 3,894 DEGs between 4 and 60 µmol photons m^−2^ s^−1^ under BL. The same pattern was observed under WL with 768 DEGs between 4 and 10 µmol photons m^−2^ s^−1^ up to 2,486 DEGs between 4 and 60 µmol photons m^−2^ s^−1^ (Fig. 4b and S7, Supporting Information Table S3). These results show that light intensity elicits a dose-dependent transcriptional response in *P. calceolata*, with a stronger effect observed under BL compared to WL. Similarly, to understand whether the observed transcriptomic regulation is steadily increasing or stable at a given light intensity, we performed a co-expression analysis. The analysis grouped the 5,217 DEGs (according to intensity or color) into 6 co-expression profiles (Fig. 4c). Half of these profiles show gene expression levels that progressively decrease as light intensity increases (profiles 2, 3 and 6), while the other half show gene expression levels that progressively increase with light intensity (profiles 1, 4 and 5). The different magnitudes of these changes and expression levels explain the presence of several clusters for each category.

The Uniform Manifold Approximation and Projection (UMAP) separated genes preferentially expressed at low irradiance from those preferentially expressed at high irradiance, indicating that irradiance-responsive genes shared relatively coherent expression profiles across BL and WL conditions (Fig. 4c). By contrast, genes associated with the BL-versus-WL contrast were more widely distributed and only partially segregated in the UMAP, suggesting that spectrum-responsive genes did not constitute a single, coherent expression class. Of the 469 genes differentially expressed between BL and WL, 259 (55%) were also identified in at least one irradiance contrast, revealing substantial overlap between the transcriptional responses to spectral quality and irradiance.

### Functional enrichment of P. calceolata genes under different lights

To identify biological functions associated with responses to irradiance, we performed Pfam-domain enrichment analyses on genes differentially expressed between the low-light condition and at least one of the higher irradiances (Fig. 5; Supporting Information Table S5). The chlorophyll a/b binding domain (PF00504), typical of light-harvesting complexes (LHC), was significantly enriched among both genes preferentially expressed at low irradiance and those preferentially expressed at high irradiance, indicating contrasting transcriptional responses among LHC-encoding genes. This observation is consistent with the dual role of LHC involved in light capture at low irradiances and photoprotection at high irradiance. LHCs were also enriched among genes more highly expressed under BL than under WL, showing that the expression of some LHC genes was associated with both irradiance and spectral quality (Supporting Information Fig. S9).

**Fig. 5.**
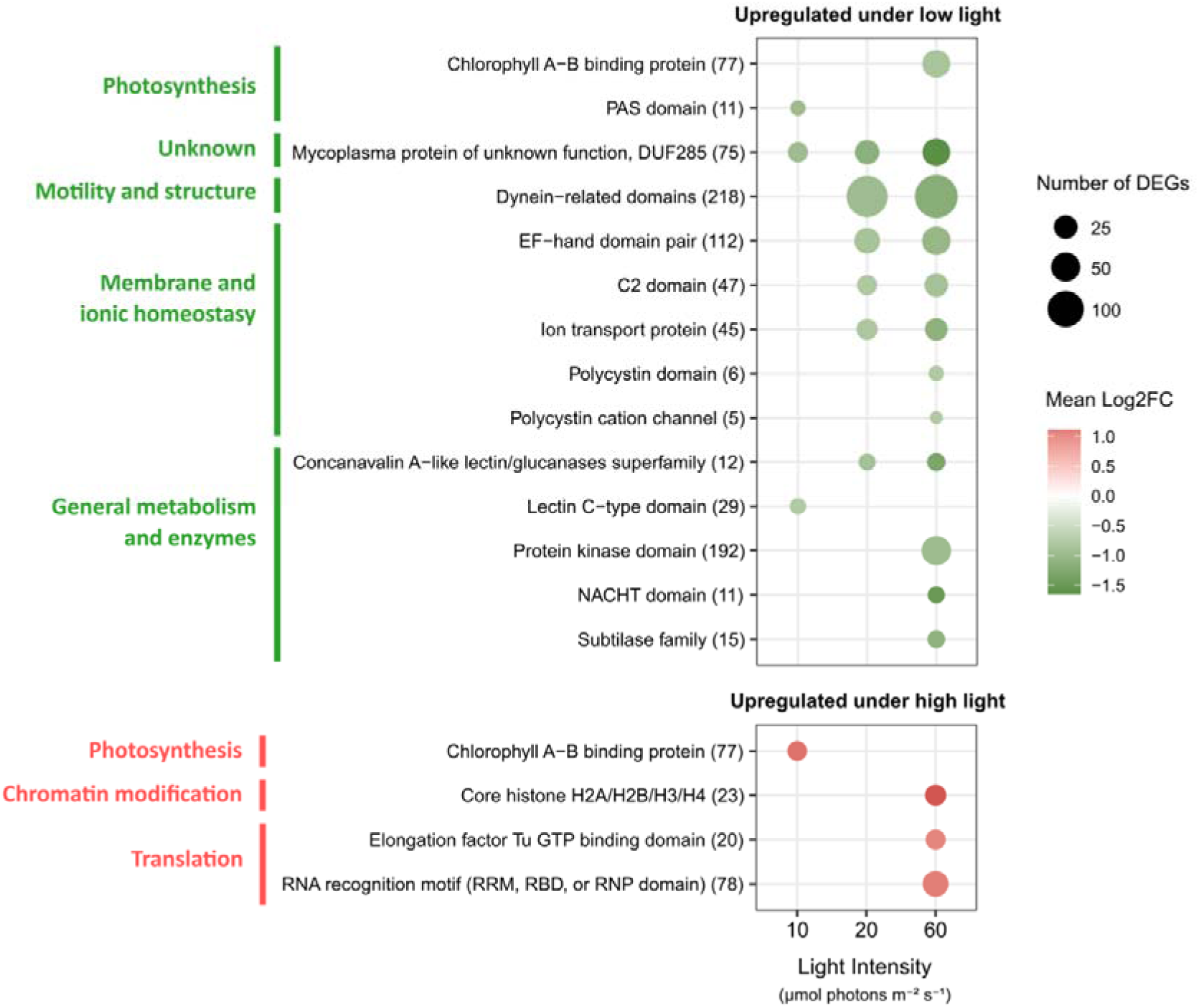
Pfam domains enrichment analysis in *P. calceolata* across light intensities. Analysis was performed on DEGs with a |log₂FC| ≥ 0.585 (corresponding to a fold change ≥ 1.5). Significantly enriched Pfam domains (hypergeometric test and Benjamini-Hochberg correction with adjusted p-value < 0.05) among DEGs upregulated under low-light (top) and under high-light (bottom) conditions are represented. Dot size indicates the number of DEGs annotated with each domain, whereas the numbers in brackets are the total number of *P. calceolata* genes annotated with each domain. Pfam IDs are provided in Supporting Information Table S5.

Genes more highly expressed under low light were enriched in several dynein-related domains associated with flagellar assembly and motility. Domains related to calcium signalling, ion or membrane transport and cellular homeostasis were also enriched, including EF-hand (PF13499), ion-transport protein (PF00520), C2 (PF00168), polycystin (PF20519) and cation-channel (PF08016) domains. Additional enriched domains included C-type lectin (PF00059), concanavalin A-like lectin/glucanase (PF13385), protein kinase (PF00069), NACHT (PF05729) and subtilase (PF00082) domains.

The domain of unknown function DUF285 (PF03382), which is abundant in microalgae and has been suggested to be involved in the cold response (Pierella Karlusich *et al*., 2025), showed progressively stronger enrichment, both in terms of number of DEGs and log₂FC, preferentially expressed at low irradiance. In the comparison between 4 and 60 µmol photons m⁻² s⁻¹, 45 of the 75 genes carrying this domain were more highly expressed at 4 µmol photons m⁻² s⁻¹. Seven of these genes were also more highly expressed under WL than under BL, indicating that a subset responded to both irradiance and spectral quality (Supporting Information Fig. S9). This result suggests that DUF285 may be involved in a broader response to environmental variations, including changes in light intensity and quality.

Among the functionally annotated genes showing the strongest response to irradiance, a gene encoding the lycopene β-cyclase (LCYB1), which catalyses the conversion of lycopene into β-carotene in the carotenoid biosynthesis pathway, displayed approximately 45-fold higher transcript abundance at 4 than at 60 µmol photons m⁻² s⁻¹ (Supporting Information Table S4). This expression pattern was consistent with the higher cellular pigment contents measured under low-light conditions.

### Modulation of photoreceptor expression levels

The large number of genes regulated by light quality and intensity suggests the presence of multiple photoreceptors capable of rapidly modulating *P. calceolata* physiology. To characterize this repertoire, we screened the genome of *P. calceolata* for genes encoding photoreceptors based on the presence of conserved functional domains and by sequence homologies with known photoreceptor proteins. A total of 29 genes encoding putative photoreceptors were identified: 7 cryptochromes/photolyases (*Cryp1-7*), including one with a CryDASH domain (*Cryp1*), 15 proteins containing one or several LOV domain(s) (*Lov1-7*), including 8 aureochromes (*Aureo1-8*), and one phytochrome-like protein (*Phyt1*) (Supporting Information Table S6). We noted that Phyt1 protein is unusually large, with a total length of 3,162 amino acids. It contains multiple PAS/PAC domains, as well as the PHY domain, characteristic of phytochromes, but no GAF domain (Supporting Information Fig. S10, Table S6). *Phyt1* is likely functionally distinct from *Phaeodactylum tricornutum* phytochrome recently identified as a detector of optical depth (Duchêne *et al*., 2025). Finally, we identified six rhodopsins based on conserved domains (*Rhod1-6*). To classified them we built a phylogeny with *P. calceolata* rhodopsins and those described in Coesel et al. 2021 (Supporting Information Fig. S9). We identified one bacteriorhodopsin (*Rhod1*), three channelrhodopsins (*Rhod2,4,6*), one sensory rhodopsin (*Rhod3*) and one heliorhodopsin (*Rhod5*).

The gene expression analysis of these 29 photoreceptors revealed heterogeneous responses to light intensity or quality, both across and within photoreceptor families (Fig. 6). Aureochromes displayed particularly variable expression patterns, with only one member (*Aureo2*) significantly upregulated (adj. p-value < 0.01, log₂FC ≥ 1) under low-light conditions, specifically under BL. The other proteins containing LOV-like domains were also more expressed under low light compared to high light for most of them, with four genes being significantly upregulated (*Lov1, Lov2, Lov4,* and *Lov7*). Only *Lov3* was significantly upregulated under high light (*Lov3*). The phytochrome-like gene (*Phyt1*) was significantly upregulated under low BL conditions. In contrast, cryptochromes were generally more expressed under high-light conditions, with two genes (*Cryp3* and *Cryp7*) showing significant upregulation. Finally, rhodopsins also exhibited diverse responses to light intensity: *Rhod1* (bacteriorhodopsin) and *Rhod3* (sensory rhodopsin) were significantly upregulated under low light, whereas *Rhod5* (heliorhodopsin) was upregulated under high light. Interestingly, the three channelrhodopsins (*Rhod2, Rhod4, Rhod6*) were showing a globally higher expression under WL compared to BL, with one (*Rhod6*) being significantly upregulated under WL, suggesting potential wavelength-specific regulation within this protein family. Collectively, these very distinct light-dependent expression patterns suggest substantial functional diversification among photoreceptors in *P. calceolata*.

**Fig. 6.**
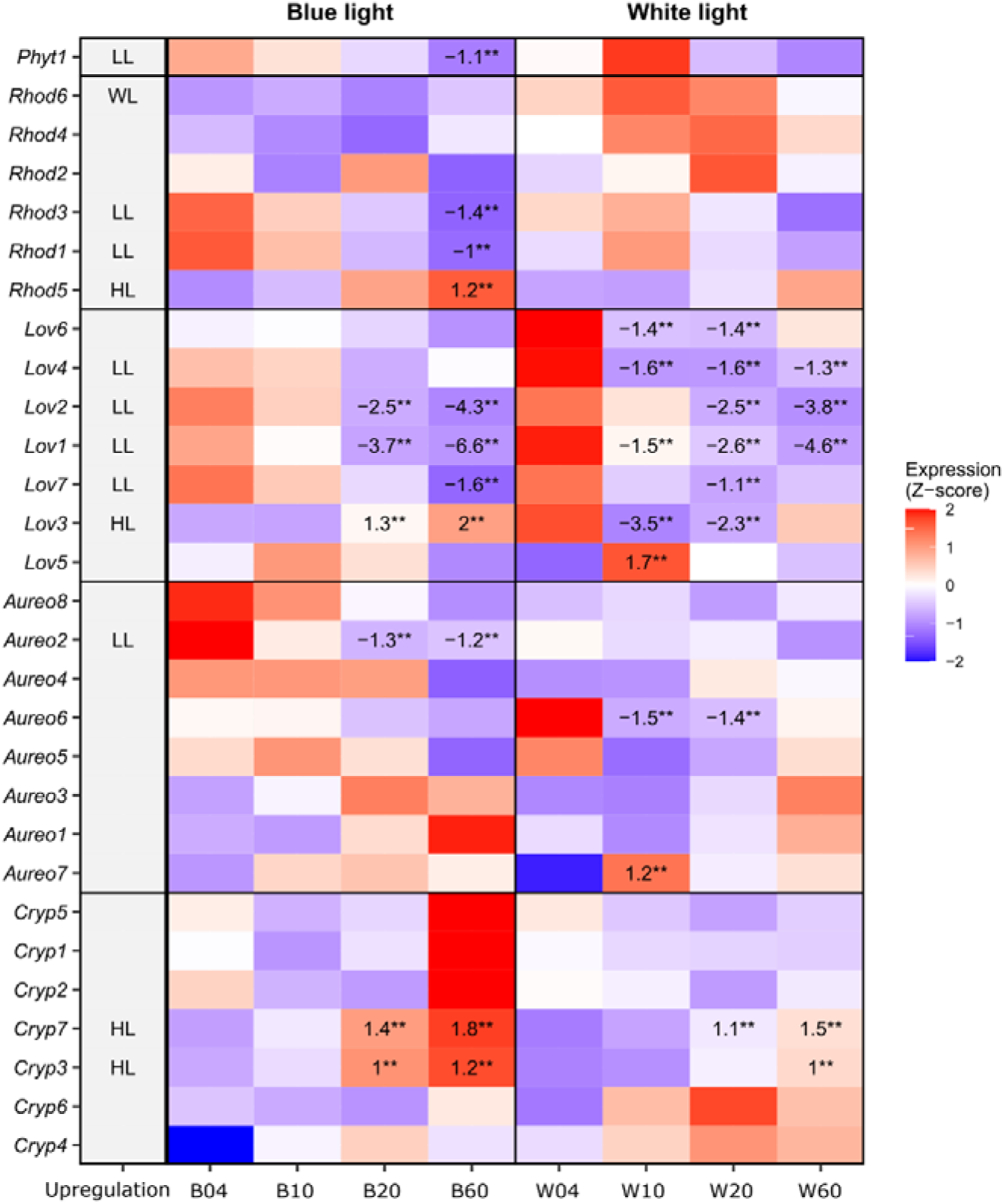
Expression profiles of photoreceptor genes at different light intensities (4, 10, 20 and 60 µmol photons m^−2^ s^−1^) under BL and WL. Normalized expression for each gene (z-score). Log₂FC relative to 4 µmol photons m^−2^ s^−1^ of BL or WL (B04 or W04) are indicated in each cell if |log₂FC| ≥ 1 and adjusted p-value < 0.01. Two stars indicate an adjusted p-value < 0.001. The “upregulation” column shows the light condition where the gene is upregulated with a |log₂FC| ≥ 1 and an adjusted p-value < 0.01. LL = low light, HL = high light, WL = white light.

### Expression of genes involved in pigment biosynthesis and photosystem stability

To determine whether the expression of genes involved in pigment biosynthesis and photosystem stability can be correlated with pigment abundance and photosynthetic efficiency in *P. calceolata*, their expression profiles were examined across the 4 light intensities under blue and white light, as well as their averaged expression under WL compared to BL. A total of 12 nuclear genes involved in the assembly and stability of photosystems were identified in *P. calceolata* genome. These genes include 3 involved in Photosystem I (PSI) and 9 involved in Photosystem II (PSII) (Fig. 7a, Supporting Information Table S7). They displayed relatively stable expression across the different light conditions. Only two were found to be significantly upregulated under high light with a log₂FC greater than 1. *Psb28_2*, known to be involved in PSII assembly and repair, particularly under light stress, and *PsbW*, known to participate in PSII dimerization and stability.

**Fig. 7.**
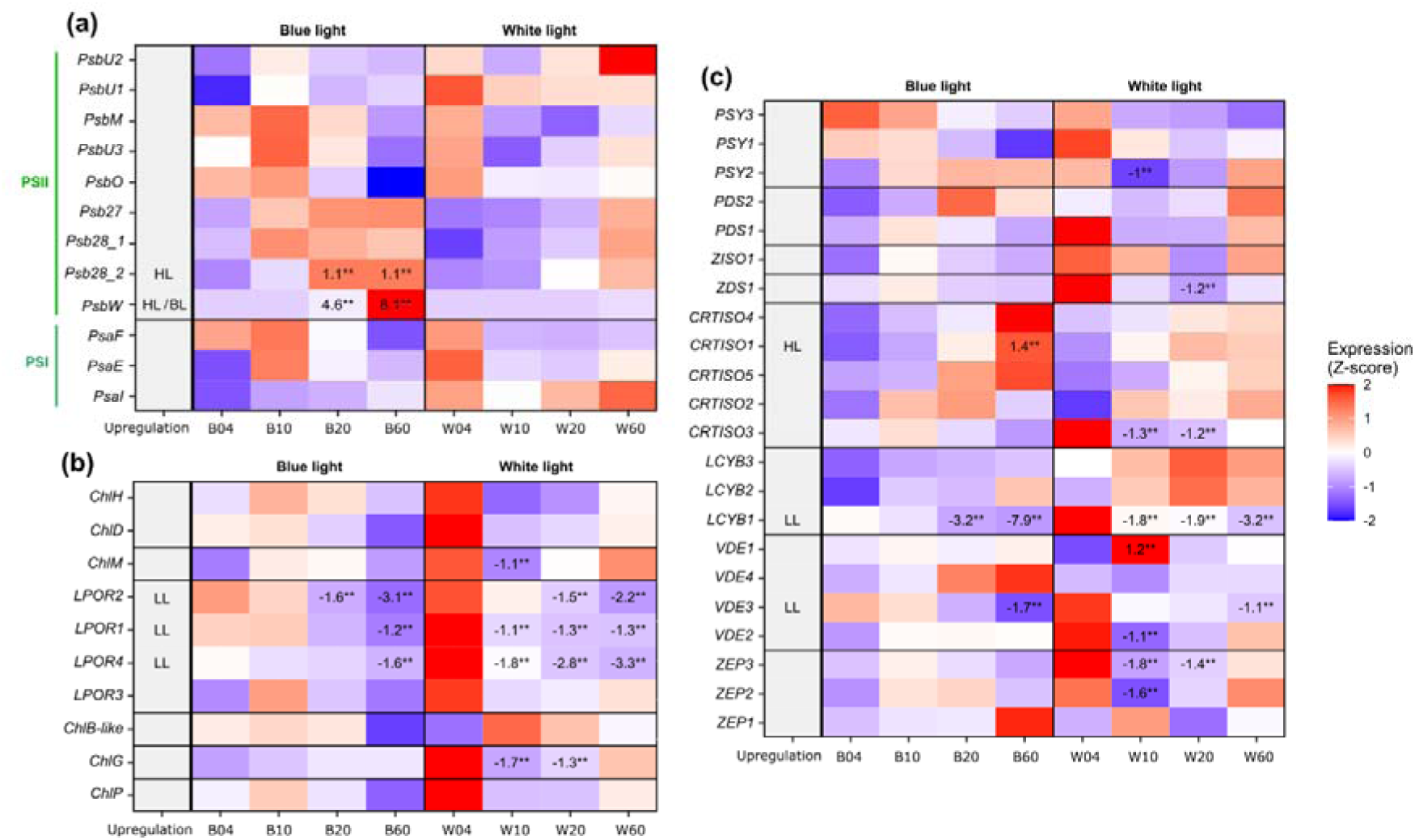
Normalized expression levels of *P. calceolata* genes involved in chlorophylls and carotenoids biosynthesis, under different light conditions. Expression of genes involved in **(a)** photosystems assembly and repair, **(b)** carotenoid biosynthesis, and **(c)** chlorophyll biosynthesis. Expression values are scaled per gene (z-score of normalized counts). Log₂ fold changes relative to 4 µmol photons m^−2^ s^−1^ of BL or WL (B04 or W04) are indicated in each cell if log₂FC ≥ 1 and adjusted p-value < 0.01. One star indicates an adjusted p-value < 0.01, and two stars an adjusted p-value < 0.001. The “upregulation” column shows the light condition where the gene is upregulated with a log₂FC ≥ 1 and an adjusted p-value < 0.01. LL = low light, HL = high light, BL = blue light.

Regarding genes involved in chlorophyll pigment biosynthesis, 25 genes were identified, including 13 involved in the tetrapyrrole common pathway, 2 encoding enzymes involved in chlorophyll modification, and 10 specific to chlorophyll biosynthesis (Supporting Information Table S7). These 10 genes are generally more expressed under low light including 3 light-dependent protochlorophyllide oxidoreductases (*LPOR1,2,4*) that were significantly upregulated in agreement with the higher abundance of chlorophyll a at low light. Interestingly, all chlorophyll pigment biosynthesis genes reached their highest expression levels under 4 µmol photons m^−2^ s^−1^ of WL, suggesting that the response is driven more by light quality than by irradiance alone (Fig. 7b).

In the *P. calceolata* genome, 22 genes are specifically involved in carotenoid biosynthesis, based on functional annotations (Fig. 7c, Supporting Information Table S7). These included 3 putative phytoene synthases (*PSY1-3*), 2 putative phytoene desaturases (*PDS1-3*), one putative zeta-carotene isomerase (*ZISO1*), one putative zeta-carotene desaturase (*ZDS1*), 5 putative prolycopene isomerases (*CRTISO1-5*), 3 putative lycopene β-cyclases (*LCYB1-3*), 4 putative violaxanthin de-epoxidases (*VDE1-4*), and 3 putative zeaxanthin epoxidases (*ZEP1-3*). Among these, the lycopene β-cyclase *LCYB1* showed the strongest differential expression under low irradiance (4 vs 60 µmol photons m⁻² s⁻¹), that mirror the abundance of the total fucoxanthin pool (Fx + 19′-BFX) under BL and WL, with a higher expression under low light, but even higher under low WL (Fig. 7c, Supporting Information Fig. S4). Similarly, the *VDE3* gene also showed a comparable pattern. In contrast, the *VDE4* gene showed an increasing expression with light intensity, only under BL, although the log₂FC was below 1. Finally, the *CRTISO1* gene, known to be involved in the isomerization of the prolycopene into lycopene, was significantely upregulated under high BL. Taken together, these results suggest that only a limited number of carotenoid biosynthetic genes are strongly regulated by light in *P. calceolata*.

## Discussion

In this study, we investigated how irradiance and spectral quality shape photoacclimation in *Pelagomonas calceolata*. We used simplified proxies for the light conditions encountered on the ocean surface, characterized by a higher irradiance and broad spectrum (white light), and the DCM, characterized by a low light intensity and reduced spectrum around blue wavelengths. Irradiance emerged as the main driver of physiological and transcriptional variation, whereas spectral quality had a more restricted effect on global gene expression but strongly influenced the 19′-butanoyloxyfucoxanthin (19′-BFx) to fucoxanthin (Fx) ratio. These results reveal a separation between broad irradiance-driven acclimation and spectrum-dependent carotenoid remodelling (Fig. 8).

**Fig. 8.**
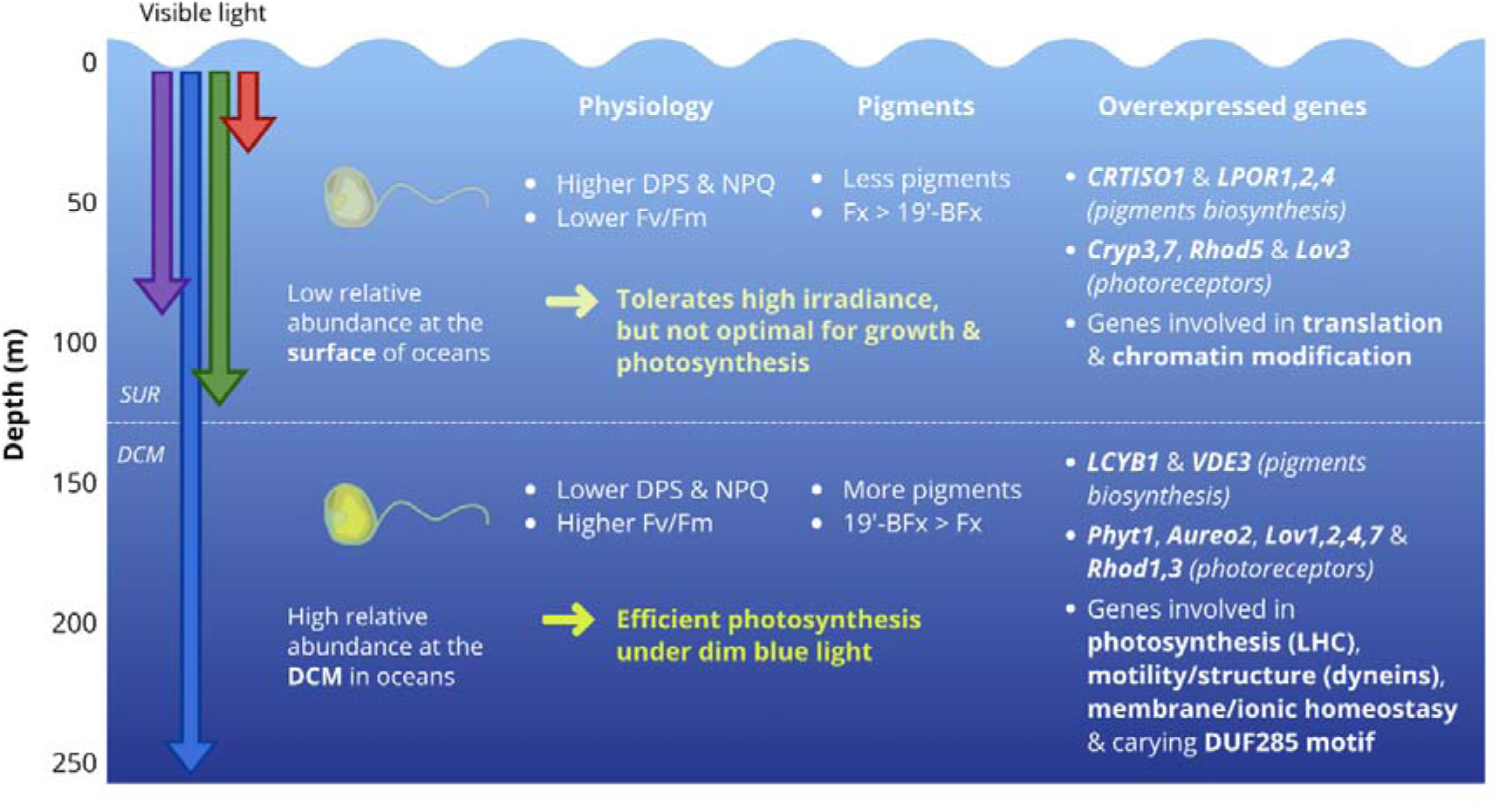
Conceptual summary of the physiological, pigment and transcriptional responses of *Pelagomonas calceolata* to irradiance and spectral quality under laboratory conditions. The penetration of the main solar wavelengths (visible range) into water is illustrated by the coloured arrows, with purple (400 nm), blue (450 nm), green (550 nm) and red (700 nm) wavelengths. The relative abundance of *P. calceolata* at the surface and at the DCM is based on previous studies (Dupont *et al*., 2015; Guérin *et al*., 2022). CRTISO = prolycopene isomerase; LPOR = light-dependent protochlorophyllide-oxidoreductase ; Cryp = cryptochrome ; Rhod = rhodopsine ; LOV = LOV-domain-containing protein ; Aureo = aureochrome ; LPOR = light-dependent protochlorophyllide oxidoreductase.

### Irradiance is the dominant factor driving photoacclimation in *P. calceolata*

Our results demonstrate that irradiance exerts a much stronger influence on the physiology and transcriptome of *P. calceolata* than spectral quality. Cells grown under low-light conditions displayed higher fluorescence and pigment contents per cell than high-light-acclimated cells, whereas comparatively little variation in total pigment content was observed between BL and WL, despite marked spectrum-dependent changes in the 19′-BFx/Fx ratio. These results are consistent with previous observations showing that changes in cellular pigment contents with increasing depth in water are primarily driven by decreasing irradiance rather than spectral composition alone (Dring, 1981; Dubinsky & Stambler, 2009). *P. calceolata* growth rate was optimal at rather dim light (10.8 µmol photons m⁻² s⁻¹ for BL, and 17.9 µmol photons m⁻² s⁻¹ for WL) (Fig. 1a) and also reached higher cell densities under low irradiance, with maximum densities observed around 10 µmol photons m⁻² s⁻¹ under both lights (Supporting Information Fig. S2). This preference for low irradiance is consistent with the ecology of *P. calceolata*, which is frequently enriched at the DCM, where photon availability strongly constrains photosynthesis (Dimier *et al*., 2009; Guérin *et al*., 2022; Coale *et al*., 2026).

Increasing irradiance triggered several physiological responses associated with photoprotection. The de-epoxidation state of the diadinoxanthin–diatoxanthin cycle increased with irradiance and broadly paralleled the increase in endpoint NPQ (Fig. 1d and 2d), while most pigment contents per cell decreased (Supporting Information Fig. S5), consistent with a reduction in cellular light-harvesting capacity. NPQ dissipates excess absorbed energy as heat and represents a widespread photoprotective mechanism in photosynthetic organisms (Goss & Lepetit, 2015; Zuo, 2025; van Amerongen & Croce, 2025). However, *P. calceolata* displayed low NPQ under the assay conditions, accompanied by declining Fv/Fm with increasing irradiance under both BL and WL (Fig. 1c). Dimier et al. (2009) similarly reported activation of the xanthophyll cycle and NPQ in *P. calceolata* RCC103 exposed to fluctuating high light, without complete protection against photoinhibition. The response to increasing irradiance was also reflected at the transcriptomic level, with seven genes encoding heat-shock proteins preferentially expressed at 60 µmol photons m⁻² s⁻¹ (Supporting Information Table S4). Overall, the low-light growth optimum and the decline in photosynthetic performance at higher irradiance are consistent with the ecological distribution of *P. calceolata* in the lower part of euphotic zone (Li *et al*., 2013).

A substantial proportion of the genes preferentially expressed under low light contained the domain of unknown function DUF285. Relative to 4 µmol photons m⁻² s⁻¹, the number of differentially expressed DUF285 genes increased at 10, 20 and 60 µmol photons m⁻² s⁻¹, while their log₂ fold changes became progressively more negative (Supporting Information Table S5). DUF285 consists of a 25-amino-acid repeat rich in leucine and lysine residues and occurs in multiple copies in the genomes of diverse microorganisms, including eukaryotic phytoplankton, bacteria and giant viruses (Röske *et al*., 2010; Moniruzzaman *et al*., 2014). DUF285 genes and transcripts are also abundant in high-latitude oceanic regions, and the predicted domain adopts a solenoid conformation resembling that of some antifreeze proteins (Pierella Karlusich *et al*., 2025). Moreover, DUF285 genes were previously found to be upregulated under severe nitrate limitation in *P. calceolata* (Guérin *et al*., 2025). Together, these observations suggest their possible involvement in a broader acclimation or stress response to several environmental fluctuations, although DUF285 function remains unknown.

*LCYB1*, encoding a lycopene β-cyclase, displayed the largest expression difference between lowest and highest irradiance among the functionally annotated genes of *P. calceolata* (log₂FC = −5.5; Supporting Information Table S4). Its expression was also influenced by spectral quality, being approximately seven-fold higher under WL than under BL (log₂FC = 2.9). Lycopene β-cyclase catalyses the cyclisation of lycopene to form β-carotene, a key branch-point reaction providing precursors for downstream xanthophyll biosynthesis (Dambek *et al*., 2012). The regulation of *LCYB* varies among taxa and physiological contexts: *LCYB* genes were little affected by high-light exposure in *Phaeodactylum tricornutum* but were induced under high light in *Haematococcus pluvialis* (Steinbrenner & Linden, 2003; Cordero *et al*., 2012; Nymark *et al*., 2013). In *P. calceolata*, neither *LCYB2* nor *LCYB3* responded significantly to irradiance, indicating differential transcriptional regulation among the three paralogues and raising the possibility that *LCYB1* has a specialised role in low-light acclimation. Similarly, most multigene families involved in carotenoid biosynthesis contained at least one gene preferentially expressed under low light, whereas other family members responded to high irradiance or showed little variation. These contrasting expression patterns are consistent with regulatory divergence following gene duplication, although they do not demonstrate biochemical subfunctionalisation. Under low light, the induction of genes such as *PSY3*, *LCYB1*, *VDE3* and *CRTISO3* may contribute to carotenoid remodelling, whereas high-light-responsive paralogues such as *CRTISO1* may participate in responses to excess irradiance (Fig. 7). Functional analyses will be required to determine whether these expression patterns redirect carotenoid metabolic flux.

Low-light acclimation was also associated with the upregulation of genes related to flagellar assembly and motility, including genes encoding dynein-related domains, as well as genes carrying calcium-binding and signalling domains such as C2 and EF-hand domains. This pattern is consistent with the calcium-mediated regulation of axonemal dynein activity and may indicate changes in flagellar activity under low-light (Smith & McIntosh, 2002; Lindemann & Lesich, 2010). Although transcript abundance alone does not demonstrate increased motility, such a response could contribute to the vertical positioning of *P. calceolata* within the water column. Similar light-dependent behavioural responses have been reported in other photosynthetic microorganisms (Cullen & MacIntyre, 1998; Clegg *et al*., 2007; Berthold *et al*., 2008), and light has been shown to have the strongest influence on phototaxis among physico-chemical parameters for most microalgae (Clegg et al. 2004).

### Candidate photoreceptors display contrasting responses to light

The *P. calceolata* genome contains a large diversity of genes encoding putative photoreceptors, suggesting the potential to sense changes across different regions of the light spectrum. Most of the identified candidates belong to blue-light-absorbing photoreceptor families, including cryptochromes and LOV-domain proteins such as aureochromes, which also carry a bZIP transcription-factor domain and are specific to Ochrophyta (Hallmann, 2025). Several aureochromes and other LOV-domain genes were more highly expressed under low light, whereas two cryptochrome genes showed higher expression under high irradiance. These opposite responses indicate that the two photoreceptor families are transcriptionally regulated by light intensity in distinct ways. Aureochromes have been implicated in blue-light sensing and acclimation, photomorphogenesis and light-dependent cell-cycle regulation in other stramenopiles (Takahashi et al. 2007; Im et al. 2024), while cryptochromes have been associated with DNA repair and photoprotection (Jaubert et al. 2022; Coesel 2024; Wu et al. 2022). Aureochromes are known to be activated photochemically, so their transcript levels reflect the size of the photoreceptor pool rather than its activation state. Their accumulation under low light may in *P. calceolata* could thus increase its ability to sense blue light, although this remains to be confirmed at the protein level.

The identification of a putative heliorhodopsin, channelrhodopsins, a sensory rhodopsin and a bacteriorhodopsin suggests an additional capacity to respond to light. Some rhodopsins regulate phototaxis in flagellated microorganisms by coupling light perception to changes in flagellar activity (Coesel *et al*., 2021). Given that *P. calceolata* is flagellated and that motility-related genes were enriched under low light, the rhodopsin candidates may participate in behavioural responses to the light environment.

Finally, the single phytochrome-like protein identified in *P. calceolata* is particularly large (3,162 amino acids), exceeding most phytochromes reported in microalgae. Its sequence shows similarity to phytochromes-like of *Skeletonema japonicum* and *A. anophagefferens*, with extensive PAS/PAC domain repetition, possibly reflecting an Ochrophyta-specific domain duplication event (Supporting Information Fig. S10, Table S6). However, the absence of a GAF domain, required for chromophore (bilin) binding, warrants caution in classifying this protein as a true phytochrome. In Ochrophyta, diatom phytochromes (DPH) are known to act as depth sensors, photoreversibly alternating between red-and far-red-absorbing forms, and to regulate gene expression in response to light (Jaubert *et al*., 2022; Hallmann, 2025; Duchêne *et al*., 2026). However, a recent study showed that the DPH enzyme of *P. tricornutum* responds to blue-green light in addition to red and far-red light, with an increased activity (Duchêne *et al*., 2025). Notably, in *P. tricornutum* and *T. pseudonana*, DPH protein levels remain constant across light conditions, including under low-blue-light regimes; it is the photoequilibrium between Pr and Pfr forms, rather than DPH abundance, that varies with the light environment (Duchêne *et al*., 2025). This contrasts with our observation in *P. calceolata*, where *Phyt1* transcript levels increase under low blue light relative to higher intensities, suggesting that the *P. calceolata* phytochrome-like protein may be subject to transcriptional regulation by light intensity, in addition to any post-translational photoconversion.

Taken together, these observations reveal that *P. calceolata* encodes a sophisticated and functionally diversified set of putative light-sensing machinery, with photoreceptor classes showing distinct, intensity- and wavelength-specific expression patterns. This result points to a highly integrated regulation of cellular responses to light quality and quantity, consistent with life in a dynamic and blue-light-dominated environment. Nevertheless, the functional assignment of these photoreceptors currently relies on domain architecture and transcriptomic data alone; protein-level characterization and gene knockout studies will be essential to confirm their roles in *P. calceolata* biology.

### Dim blue light induces reversible 19′-BFx/Fx remodelling

We show that the 19′-BFx/Fx ratio specifically increases under dim BL in *P. calceolata*. An increase of 19′-BFx abundance under low light conditions has also been observed in the haptophyte *Phaeocystis antarctica*, in which the authors suggested a light harvesting role for 19′-BFx, although this pigment is not abundant in this species (Van Leeuwe *et al*., 2014). Interestingly, the opposite response has been observed in the pelagophyte *A. anophagefferens*, where the 19′-BFx/Fx ratio increased under high light conditions (Alami *et al*., 2012; Cui *et al*., 2025), suggesting species- or condition-dependent regulation of 19′-BFx accumulation.

In our experiments, the 19′-BFx/Fx ratio did not exhibit a clear trend across WL intensities, whereas it increased markedly under dim BL. During the spectral-shift experiment, the ratio increased gradually after transfer from WL to BL, with 19′-BFx becoming more abundant than Fx after approximately 9 days. Conversely, transferring BL-acclimated cultures back to WL initiated a decrease in the ratio that was detectable within 6 h, without significantly affecting cell growth (Fig. 3). This reversible response at constant irradiance supports regulated, spectrum-dependent pigment remodelling rather than a simple effect of culture age or growth phase. However, its asymmetric kinetics - a gradual accumulation over several days under BL but a detectable decline within hours after returning to WL - distinguish it from rapid photoprotective responses such as NPQ and instead suggest a longer-term form of chromatic acclimation. The biochemical basis of this remodelling remains unresolved, including whether it involves direct enzymatic acylation of Fx, differential synthesis of the two pigments, or differences in their turnover. Previous work in *A. anophagefferens* also suggested that regulation of the 19′-BFx/Fx ratio differs from the rapidly reversible diadinoxanthin–diatoxanthin cycle, since it is not linked to pH changes (Cui *et al*., 2025). Whether the same regulatory basis applies to *P. calceolata* remains unknown.

Physicochemical studies using ultrafast transient absorption spectroscopy have shown that the non-conjugated acyloxy group of 19′-BFx modifies its excited-state dynamics relative to Fx (Staleva-Musto et al., 2018). Studies of 19′-HFx have also shown that acylation modifies excitation-energy-transfer kinetics while maintaining a high overall transfer efficiency (Staleva-Musto *et al*., 2019). These observations raise the hypothesis that 19′-BFx accumulation could affect light harvesting under photon-limited conditions. However, our results do not demonstrate that 19′-BFx improves energy transfer or photosynthetic efficiency in *P. calceolata*. This will require determining whether 19′-BFx is incorporated into specific antenna complexes and directly measuring its contribution to energy transfer. The spectrum-dependent plasticity of the 19′-BFx/Fx ratio observed here may also affect the interpretation of 19′-BFx as a chemotaxonomic marker. Although this pigment remains informative when used alongside other diagnostic pigments to identify pelagophyte-containing communities, variation in its cellular abundance means that its environmental concentration may not scale directly with pelagophyte biomass. Consequently, changes in 19′-BFx concentration may reflect both variations in community composition and physiological acclimation within 19′-BFx-producing populations.

Overall, irradiance was the dominant driver of physiological and transcriptional acclimation in *P. calceolata*, whereas spectral quality elicited a more targeted response. Within this spectral response, dim blue light promoted a gradual and reversible increase in the 19′-BFx/Fx ratio, accompanied by colour-dependent transcriptional changes. These findings identify 19′-BFx remodelling as a component of long-term chromatic acclimation while emphasizing that photon availability remains the primary determinant of photoacclimation in this low-light-adapted pelagophyte.

## Supporting information

SuppInfo

## Data availability statement

The raw RNA-sequencing reads generated in this study are available in the European Nucleotide Archive under project accession PRJEB124428 (Supporting Information Table S8). *Pelagomonas* genome sequences and gene models are available under ENA project PRJEB48576.

## Acknowledgements

We thank the following people and institutions whose commitment made this work possible: the Genoscope/CEA, Paris-Saclay University and the CNRS; the members of the Roscoff Culture Collection for providing the RCC100 strain; Carine Vergne for her assistance with the development of the HPLC protocol; and Claude Scarpelli for his support with high-performance computing at Genoscope. We gratefully acknowledge financial support from the ANR (ANR-22-CE20-0012).

## Author contributions

CS, AT and QC conceived and planned this study. CS performed *P. calceolata* cultures and pigment analyses, with support from AT, CO, LB and AJF. CS performed the RNA extractions, and AM coordinated library preparation and sequencing at the platform. CS carried out the bioinformatic analyses, supervised by QC. BN performed the second round of gene annotation for RCC100, with support from QC and CS. CS, AT and QC wrote the manuscript. QC secured funding for the project. All authors contributed to manuscript preparation and approved the final version of the paper.

---

**Supporting Information Fig. S1.**
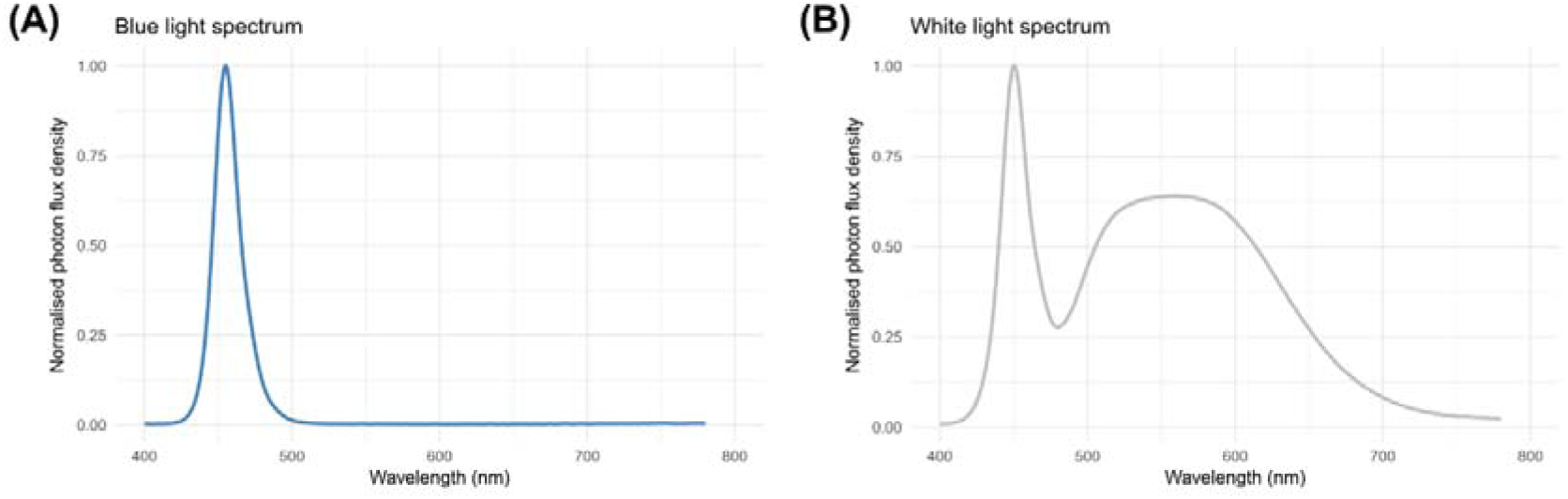
Blue and white light emission spectra of LEDs used in this study. PFD (photon flux density) values are normalised.

**Supporting Information Fig. S2.**
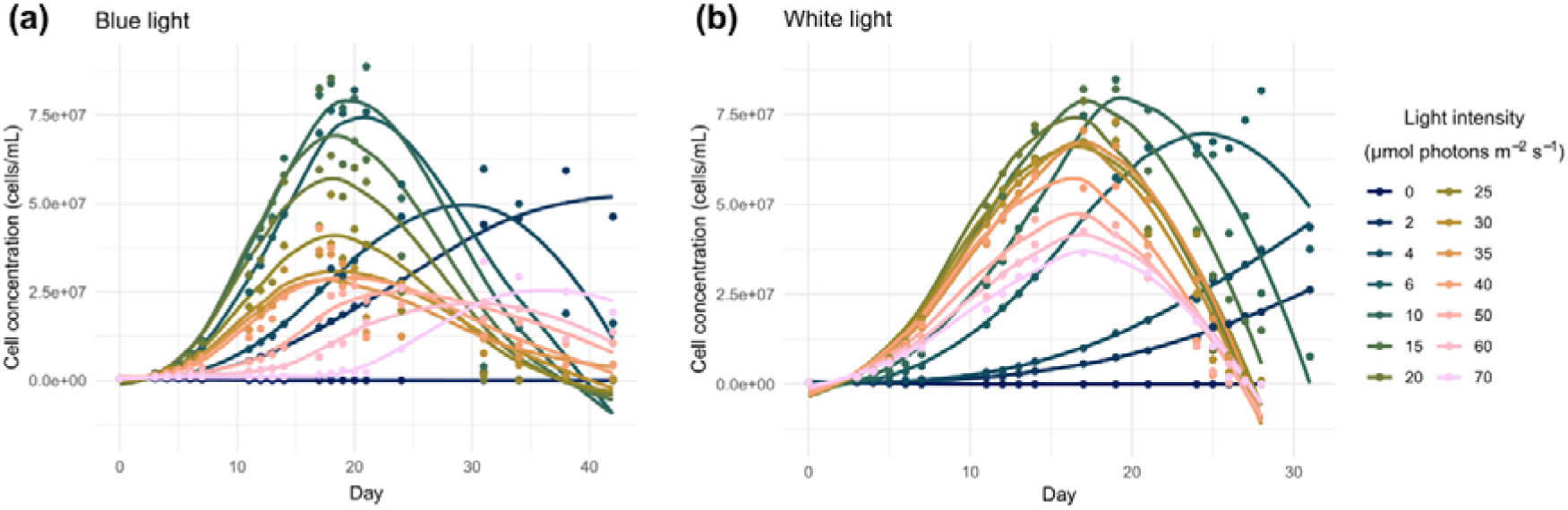
*P. calceolata* cell concentration (cell/mL) during 42 days of culture under 14 intensities of blue light (left), and during 31 days of culture under 15 intensities of white light (right). Regression lines were traced using *loess* function with R package ggplot2.

**Supporting Information Fig. S3.**
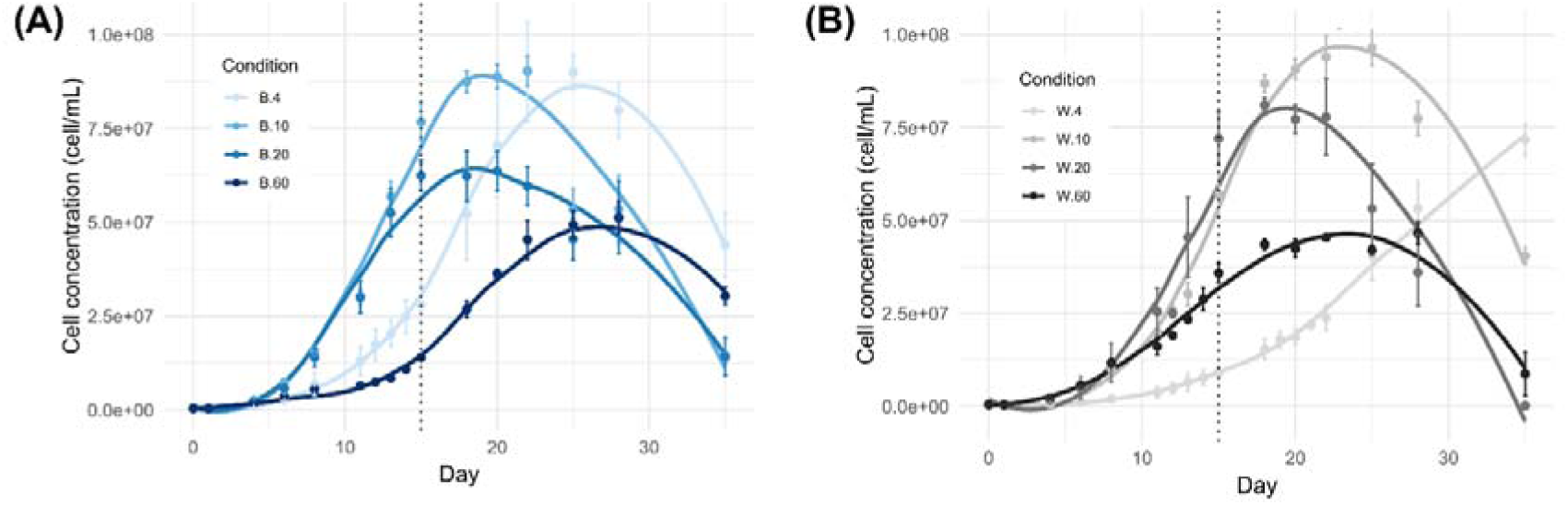
Evolution of *P. calceolata* cell density (cell mL^−1^) during 35 days of culture under 4 intensities of blue (A) and white (B) lights. The vertical dotted lines correspond to the harvesting day for the pigments analysis.

**Supporting Information Fig. S4.**
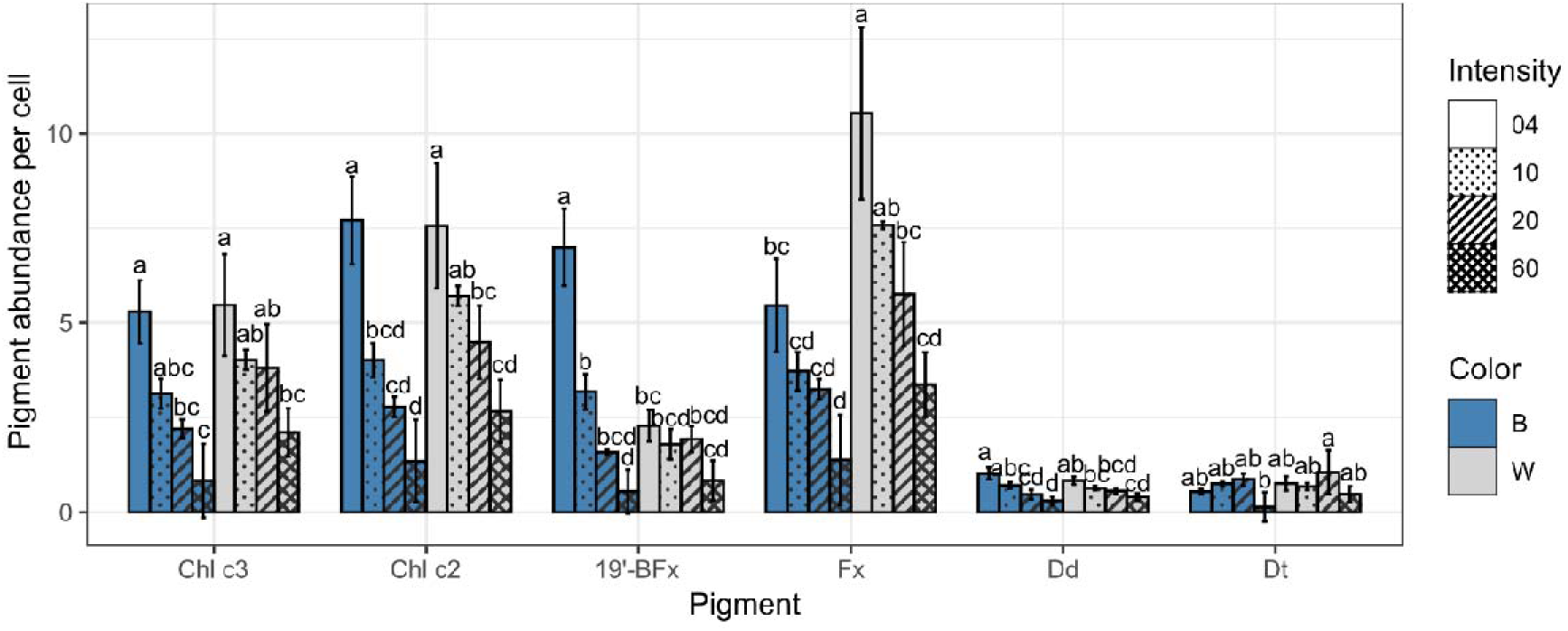
Pigment abundances per *P. calceolata* cell under different intensities of blue light (B) and white light (W). Pigment abundances are divided by the number of cells estiomated by flow cytometry.

**Supporting Information Fig. S5.**
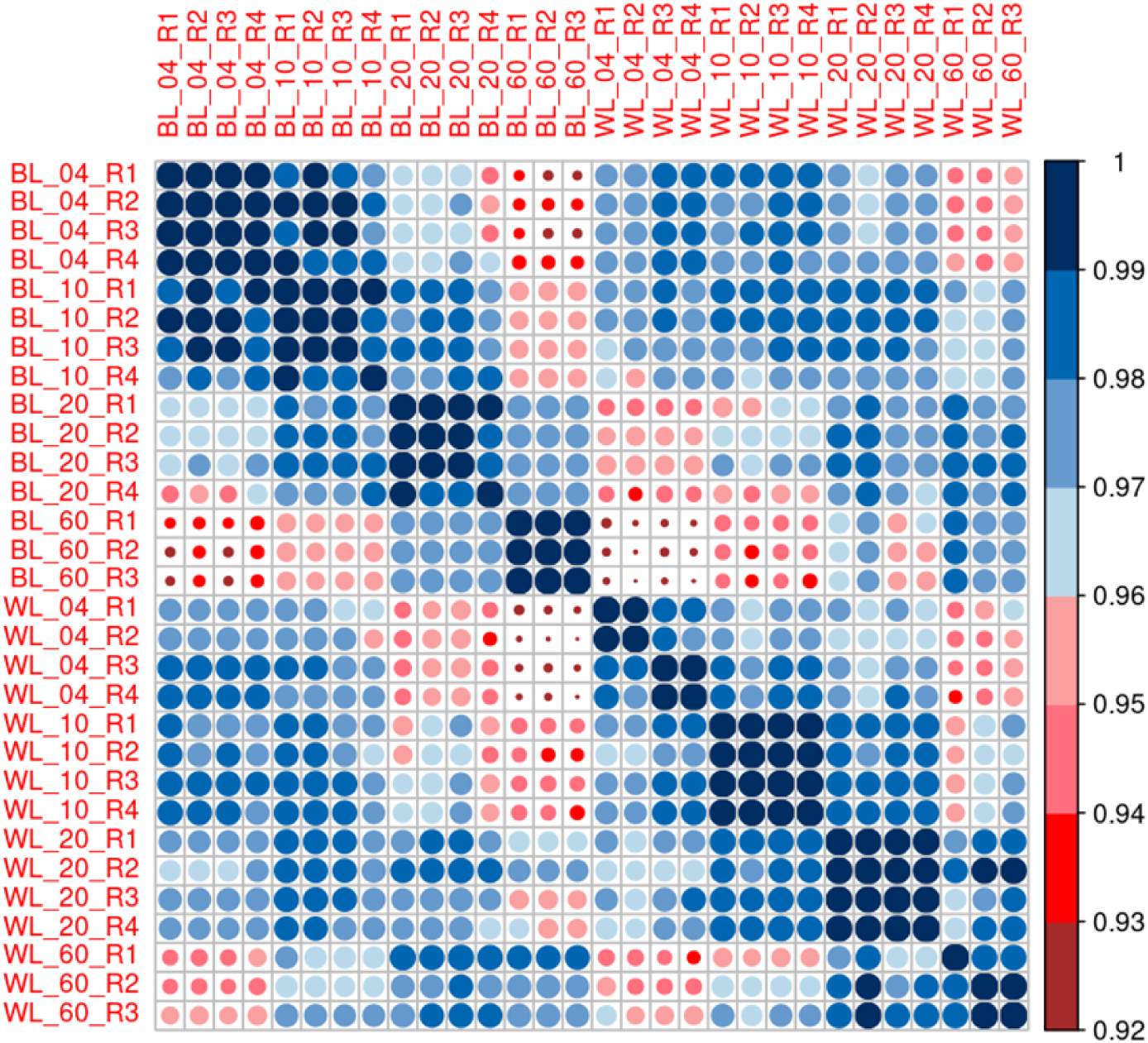
Pearson correlation matrix of the 30 *P. calceolata* samples grown under four light intensities of blue (BL) and white (WL) light. Correlations were computed on variance-stabilised counts (vst).

**Supporting Information Fig. S6.**
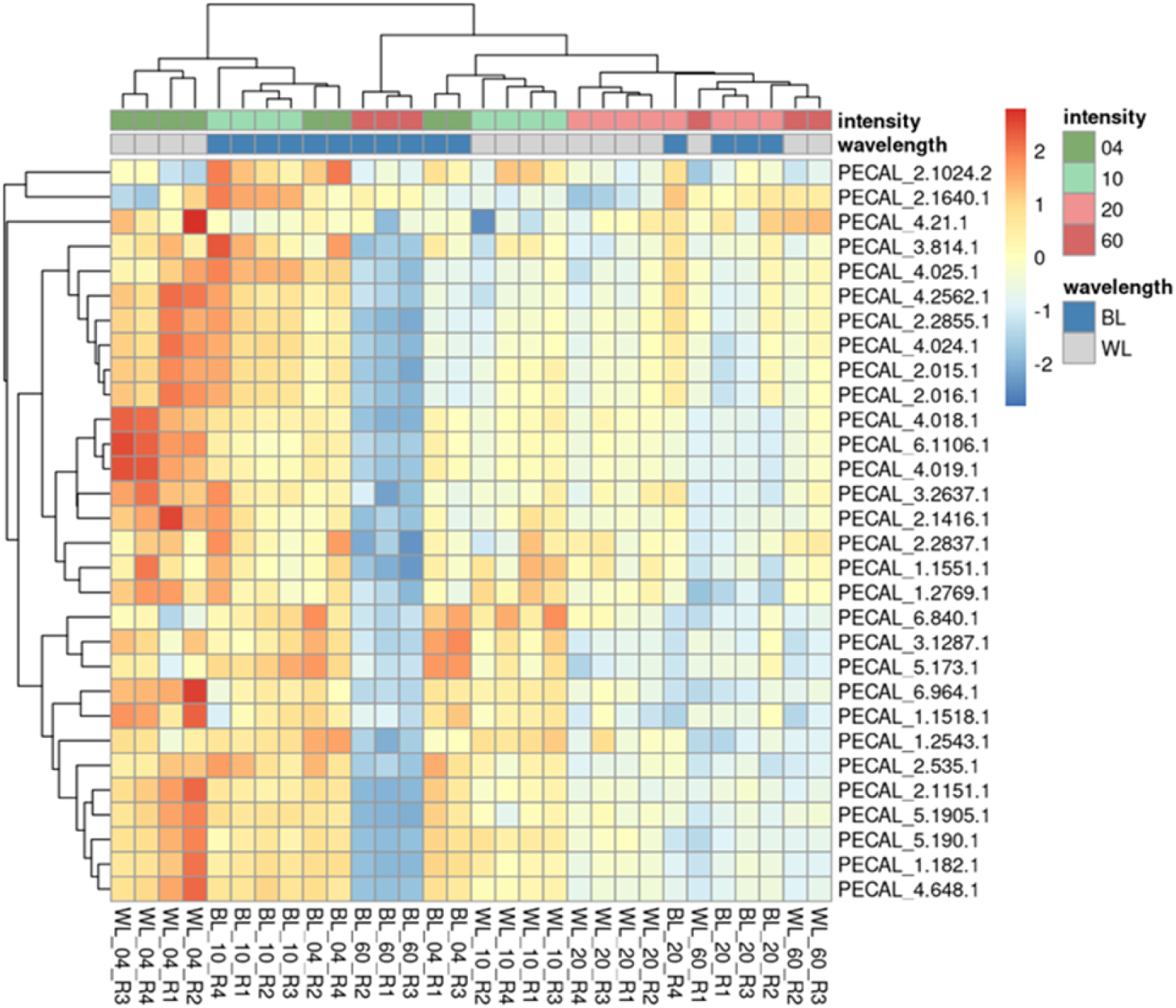
Heatmap of normalized gene expression of the 30 genes with the highest variance. The upper clustering is grouping the samples with similar expression patterns across genes, while the left clustering is grouping the genes with similar counts across samples. The gene expression is normalized by TPM method, and by line (Z-score).

**Supporting Information Fig. S7.**
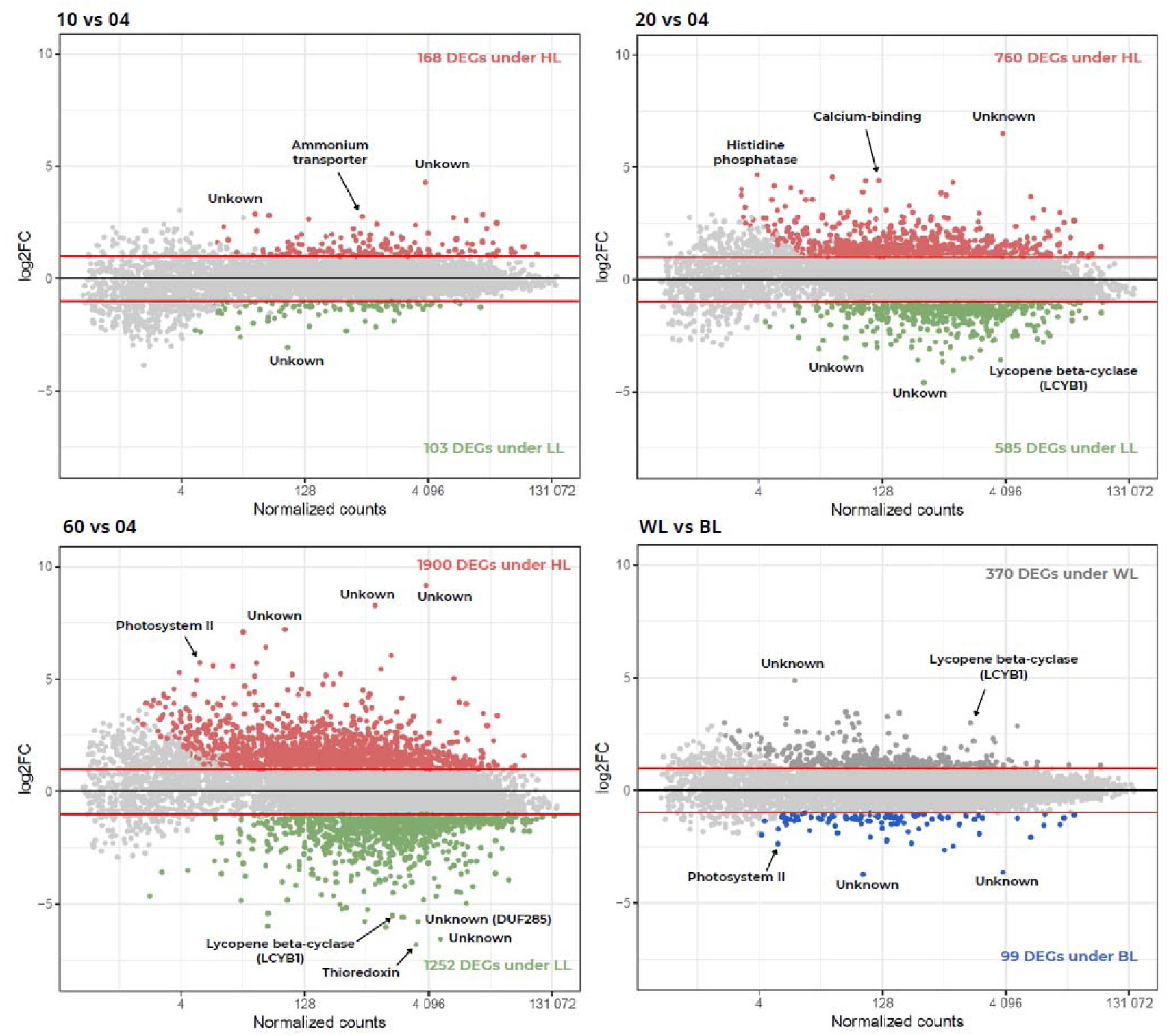
Log₂FC and counts of DEGs between the different light intensities (BL and WL). MA-plots showing the counts (x axis) and the log₂FC (y axis) of the DEGs (adjusted p-value < 0.01 and |log₂FC| > 1) under 10, 20, and 60 µmol photons m⁻² s⁻¹ compared with the 4 µmol photons m⁻² s⁻¹ condition, under both BL and WL.

**Supporting Information Fig. S8.**
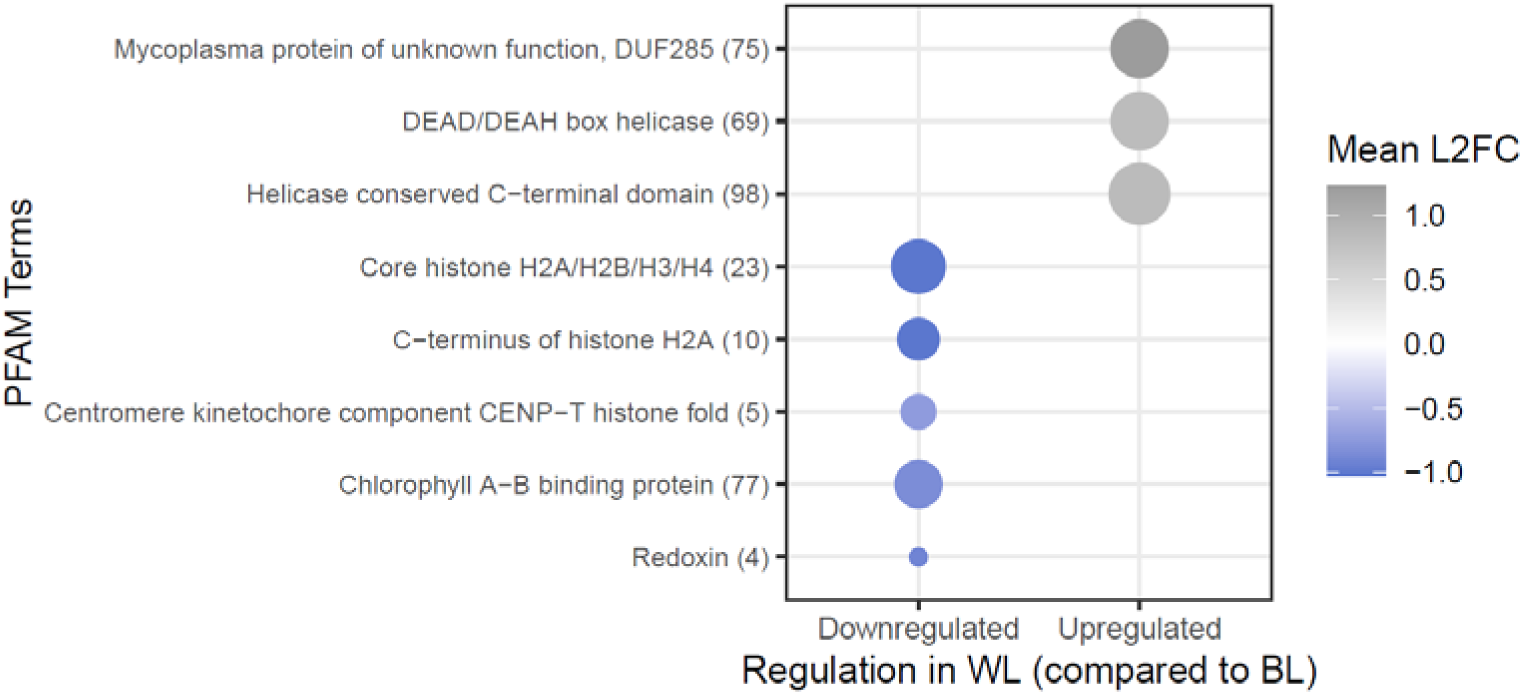
Pfam enrichment in *P. calceolata* cells under white light (WL) compared to blue light (BL). Analysis was done on DEGs with a |log₂FC| ≥ 0.585 (fold-change ≥ 1.5). Significantly enriched Pfam domains (hypergeometric test and Benjamini-Hochberg correction with adjusted p-value < 0.05) among DEGs upregulated under low light (top) and under high light (bottom) are represented. The dot size represents the number of DEGs annotated with each domain, whereas the numbers in brackets are the total number of P. calceolata genes annotated with each domain.

**Supporting Information Fig. S9.**
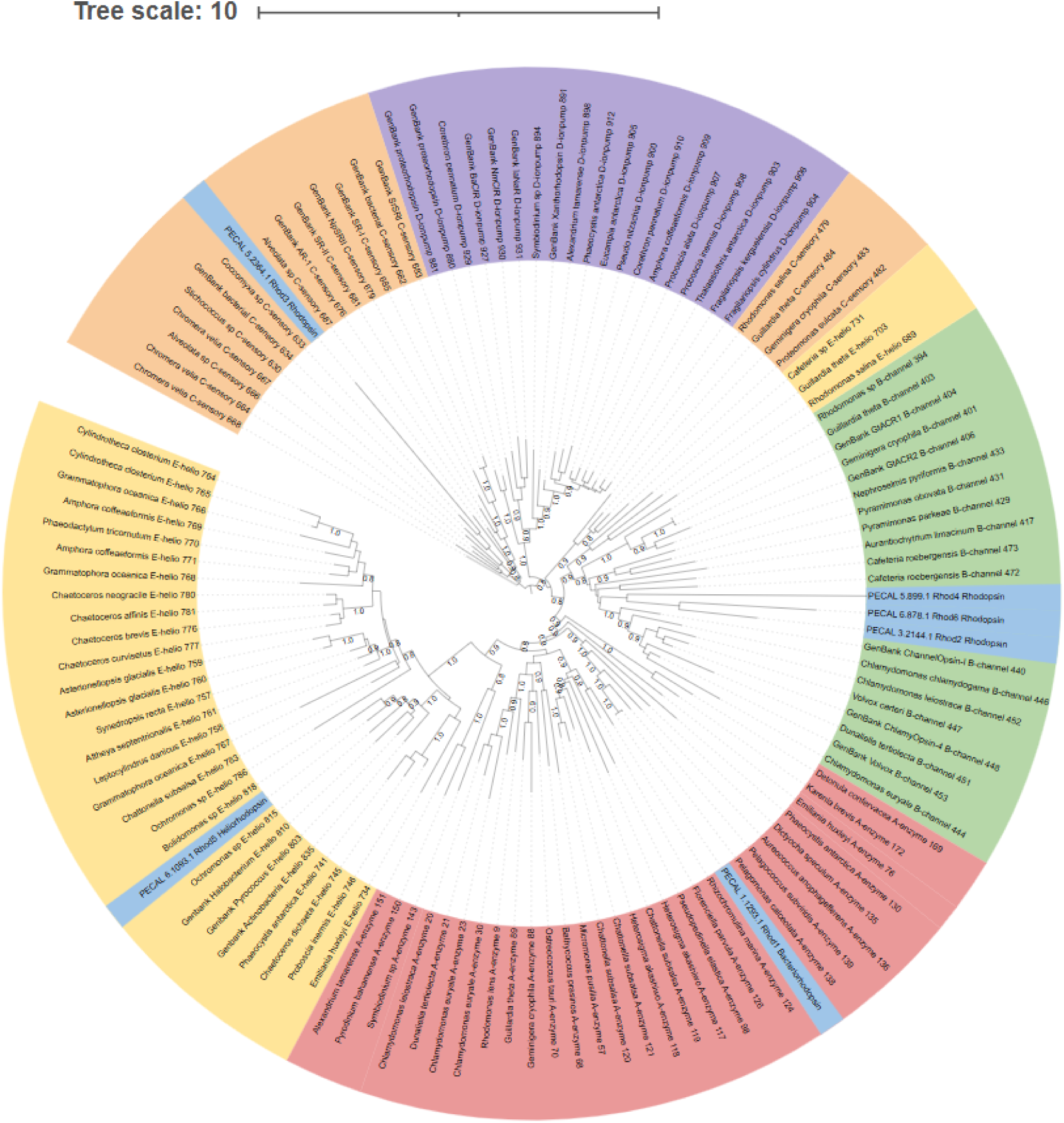
Phylogeny of microalgae rhodopsins. The tree includes the 6 rhodopsin proteins of *P. calceolata* (colored in blue) and a subset of 116 rhodopsin proteins from different groups (enzyme rhodopsins, channelrhodopsins, sensory rhodopsins, ion-pump rhodopsins and heliorhodpsins) from Coesel et al., 2021. Branch support: Transfer Bootstrap Expectation (TBE) from 100 bootstrap replicates.

**Supporting Information Fig. S10.**
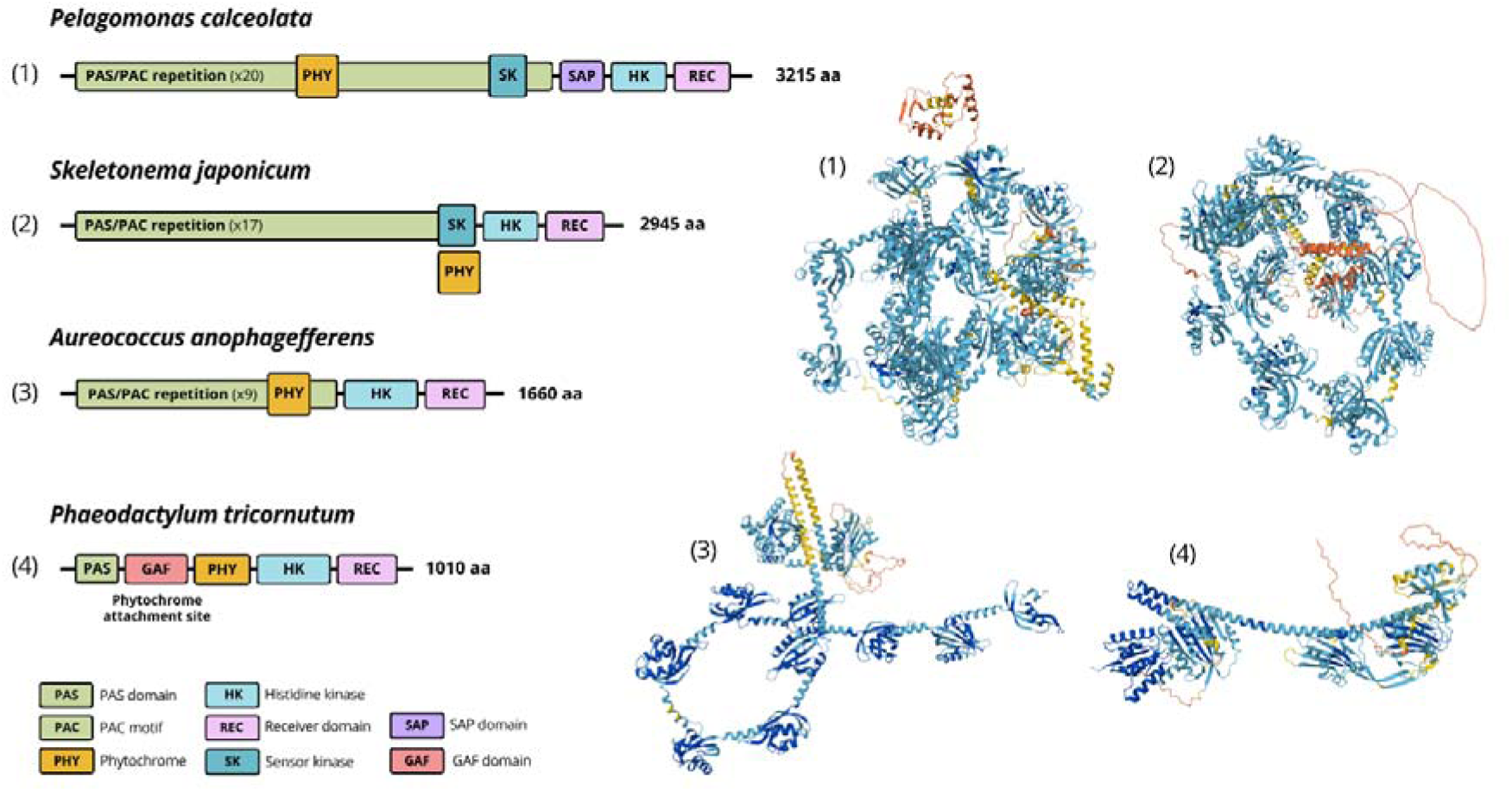
Functional domains and 3D structures of phytochromes and phytochromes-like proteins in 4 microalgae. IPR functional domains were determined using InterProScan. PAS = Per-ARNT-Sim domain (IPR000014), PAC = PAC motif (IPR001610), PHY = Phytochrome specific domain (IPR001294), SK = Sensor kinase/photoreceptor (IPR052162), HK = Histidine kinase domain (IPR005467), REC = Receiver domain (IPR001789), SAP = SAF-A/B, Acinus, and PIAS domain (IPR003034), GAF = cGMP-specific phosphodiesterases, adenylyl cyclases and formate hydrogen lyase domain (IPR003018). Number of amino acids per sequence are indicated at the end of the sequence. Structures were generated using AlphaFold 3 and colored according to the per-residue confidence metric (pLDDT): dark blue for very high confidence (predicted Local Distance Difference Test (pLDDT) > 90), light blue for high confidence (90 ≥ pLDDT > 70), yellow for low confidence (70 ≥ pLDDT > 50), and orange for very low confidence (pLDDT ≤ 50).

